# An NLP-inherited motif confers broad DNA-binding specificity to NIN in root nodule symbiosis

**DOI:** 10.1101/2025.07.20.665785

**Authors:** Shohei Nosaki, Momona Noda, Hiroki Onoda, Momoyo Ito, Takuya Suzaki

## Abstract

Nitrogen-fixing root nodule symbiosis (RNS) occurs in some eudicots, including legumes, and is regulated by the transcription factor NODULE INCEPTION (NIN), derived from the NIN-LIKE PROTEIN (NLP) family. However, how the NIN protein acquired RNS-specific functions remains unclear. We identify a previously undescribed motif in *Lotus japonicus* NIN, located downstream of the RWP-RK domain, which we term the FR. This motif broadens the DNA-binding specificity of NIN by stabilizing the RWP-RK dimer interface. *nin* mutants lacking the FR motif show defective nodulation and impaired nitrogen fixation. *Arabidopsis* NLP2 carries a NIN-type FR and shares key features with NIN. Furthermore, the NIN-type FR likely originated as early as gymnosperms, suggesting that the molecular feature of NIN for RNS regulation was inherited from ancestral NLPs before RNS emerged.

## Main Text

Root nodule symbiosis (RNS) is a symbiotic relationship between nitrogen-fixing clade plants, including legumes, and nitrogen-fixing bacteria such as rhizobia or Frankia (*1, 2*). NODULE INCEPTION (NIN) transcription factor (TF) is a master regulator of RNS, controlling rhizobial infection, root nodule development, and symbiotic nitrogen fixation by regulating key RNS-related genes, including *NUCLEAR FACTOR-YA* (*NF-YA*), *NF-YB*, *EXOPOLYSACCHARIDE RECEPTOR 3* (*EPR3*), and *RHIZOBIUM-DIRECTED POLAR GROWTH* (*RPG*) (*3–6*). Previous phylogenomic studies have suggested that the presence or absence of NIN in plants is closely linked to the acquisition and loss of RNS ability (*7, 8*). In *Medicago truncatula*, NIN undergoes protein processing during nodule development, with the C-terminal region playing a role in regulating further RNS processes (*9*). While our understanding of NIN’s function is growing, the fundamental question of which molecular features of NIN enable its regulation of RNS remains unresolved.

NIN is derived from the NIN-LIKE PROTEIN (NLP) TF family, which includes an RWP-RK DNA-binding domain and a PB1 domain involved in multimerization (*10, 11*). A GAF-like domain in the N-terminal region, responsible for nitrate sensing, is conserved among NLPs but absent in NIN (*12, 13*), which may account for NIN’s loss of nitrate responsiveness. In *Lotus japonicus*, LjNIN and LjNLP4 form homodimers and bind to similar *cis*-elements containing two palindromic core-sites on the promoters of RNS and/or nitrate-related genes (*14*) (Fig. 1, A and B). LjNIN can bind to the LjNLP4-binding sites, but LjNLP4 does not necessarily bind to LjNIN-specific sites. The *cis*-element on Pro*LjCLE-RS2* is highly palindromic and serves as a common binding site for both LjNIN and LjNLP4. In contrast, some LjNIN-specific binding sites (e.g., Pro*LjNF-YA*, Pro*LjNF-YB*) are less palindromic compared to LjNLP4-binding sites, suggesting that LjNIN has a broader DNA-binding specificity than LjNLP4 (*14*). However, the underlying mechanisms remain unclear.

**Fig. 1.**
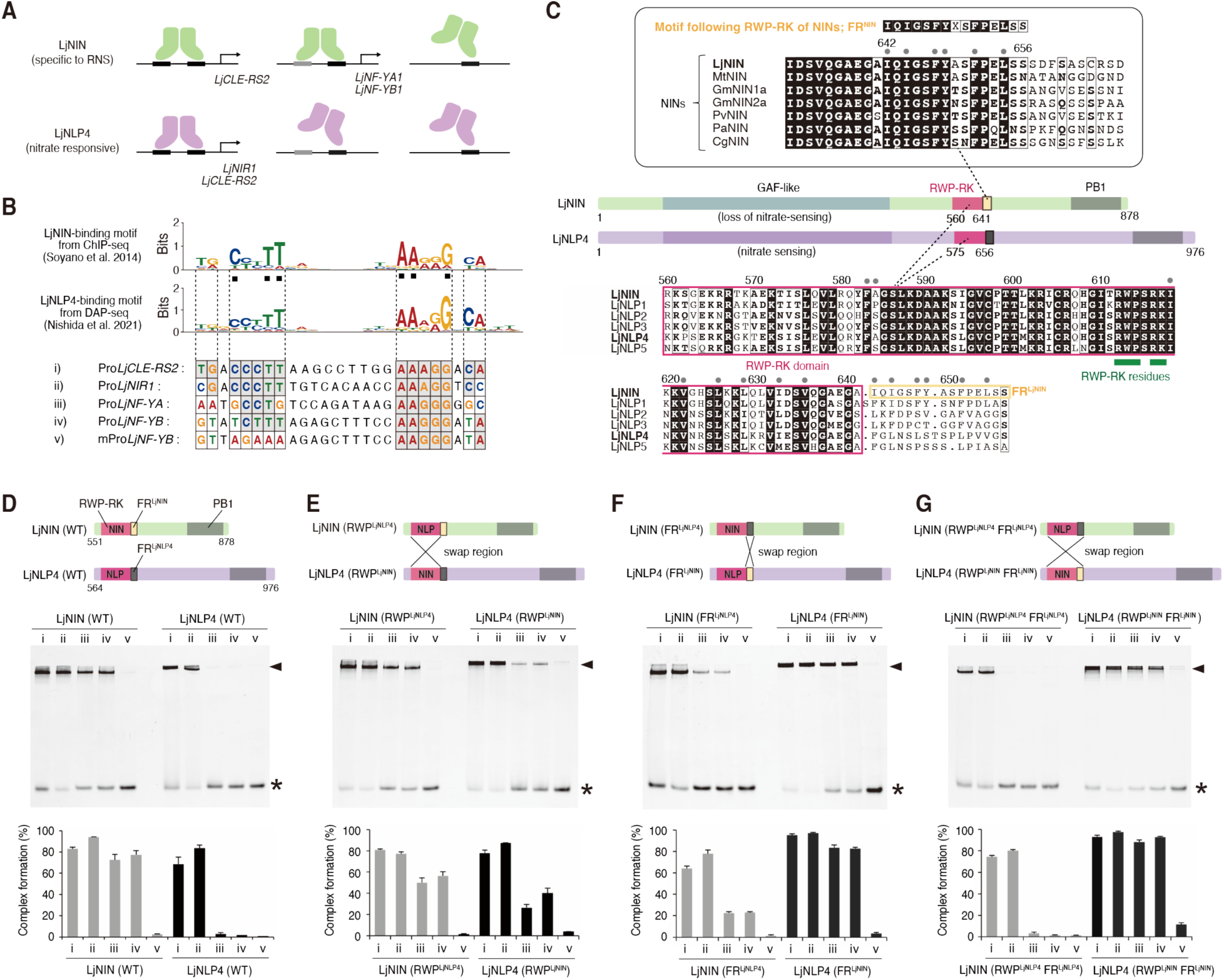
An amino acid motif that underlies the NIN’s broad DNA binding selectivity. (**A**) The broad DNA-binding selectivity of LjNIN from *Lotus japonicus* (specific to RNS) distinct from LjNLP4 (nitrate-responsible). Both LjNIN and LjNLP4 bind to perfect preferred *cis*-elements comprising two palindromic core-sites (black boxes), whereas only LjNIN can bind partially unpreferred *cis*-elements in which one of the two core-sites is partially unpreferred (gray box), which underlies the NIN-specific gene regulation. (**B**) Sequences of the fluorescence-labeled DNA probes for EMSA designed based on the LjNIN/LjNLP4 binding-motifs derived from the previously reported genome-wide analyses (*14*, *32*). (**C**) Domain structures of LjNIN and LjNLP4 with protein sequence alignments of the Motif Following RWP-RK (FR) of NINs (FR^NIN^) from *L. japonicus* (Lj), *Medicago truncatula* (Mt), *Glycine max* (Gm), *Phaseolus vulgaris* (Pv), *Parasponia andersonii* (Pa), and *Casuarina glauca* (Cg), and the RWP-RK domains with the FRs from LjNIN/NLPs. White boldface on black background and black boldfaces on white background show the completely and partially conserved amino-acid residues, respectively. The amino acid sequence identity within the RWP-RK domain among LjNIN and LjNLPs is 52%. Gray circles on alignments indicate the amino-acid residues forming a dimer interface via hydrophobic and/or van der Waals interactions in the AlphaFold2-predicted structure of the LjNIN dimer. (**D** to **G**) EMSA results of LjNIN-WT, LjNLP4-WT (D) and the LjNIN/LjNLP4 chimeras with the RWP-RK regions swapped (LjNIN (RWP^LjNLP4^) /LjNLP4 (RWP^LjNIN^); E), the FR swapped (LjNIN (FR^LjNLP4^) /LjNLP4 (FR^LjNIN^); F), and both the RWP-RK and the FR swapped (LjNIN (RWP^LjNLP4^ FR^LjNLP4^) /LjNLP4 (RWP^LjNIN^ FR^LjNIN^); G). The constructs of LjNIN/LjNLP proteins are shown at upper panels. Each recombinant protein fused to the MBP at N-terminus was reacted at 2 μM final conc. with 0.25 μM DNA probe (i to v). The electrophoretic patterns are shown at middle panels with asterisks and arrowheads indicating the positions of free DNA and the protein-DNA complexes, respectively. Bar graphs at lower panels showing the fluorescence densitometric profile are presented as mean ± s.e.m. (n=3).

## Results

### Identification of the motif determining the broad DNA-binding specificity of NIN

To identify the key amino-acid residues determining the NIN-specific broad DNA-binding specificity, we examined amino-acid conservation in NINs/NLPs across various plant species. Phylogenetic analysis revealed that the RWP-RK domains derived from NINs showed a pattern similar to that of full-length NINs (fig. S1, A and B). Amino-acid sequence alignments further showed RWP-RK-comprising residues are more conserved in NINs compared to NLPs (Fig. 1C and fig. S1C). To test whether the RWP-RK domain of NIN is crucial for its broad DNA-binding specificity, we conducted an electrophoretic mobility shift assay (EMSA) using chimeric proteins (LjNIN/LjNLP4) with swapped RWP-RK domains (LjNIN (RWP^LjNLP4^) /LjNLP4 (RWP^LjNIN^)), along with wild-type proteins (LjNIN (WT)/LjNLP4 (WT)) (Fig. 1, D and E and fig. S2). LjNIN (WT) bound strongly to the partially unpreferred *cis*-elements Pro*LjNF-YA*(iii)/Pro*LjNF-YB*(iv) and the perfectly preferred *cis*-elements Pro*LjCLE-RS2*(i)/Pro*LjNIR1*(ii), but not to the mPro*LjNF-YB*(v) with only one core-site (Fig. 1D). In contrast, LjNLP4 (WT) specifically bound to Pro*LjCLE-RS2*(i)/Pro*LjNIR1*(ii) (Fig. 1D). Under the same conditions, LjNIN (RWP^LjNLP4^) still interacted with Pro*LjNF-YA*(iii)/Pro*LjNF-YB*(iv), albeit with reduced binding compared to LjNIN (WT). LjNLP4 (RWP^LjNIN^), unlike LjNLP4 (WT), could bind to these two *cis*-elements, but less effectively than LjNIN (RWP^LjNLP4^) (Fig. 1E). These results suggest that the RWP-RK domains contribute partially, but not critically, to the distinctive DNA-binding specificities of LjNIN and LjNLP4.

We then focused on a 15-amino-acid motif located just following the RWP-RK domains, which we hereby named the Motif following RWP-RK (FR). Notably, this FR motifs in NINs (FR^NIN^) are evolutionary conserved among nodulating plants of nitrogen-fixing clade, including legumes, *Parasponia*, and an actinorhizal plant (*15, 16*) (Fig. 1C). EMSA using chimeric proteins with the FR swapped (LjNIN (FR^LjNLP4^) /LjNLP4 (FR^LjNIN^)) (Fig. 1F and fig. S2, C and D) showed that LjNIN (FR^LjNLP4^) had significantly reduced binding to Pro*LjNF-YA*(iii)/Pro*LjNF-YB*(iv) compared to LjNIN (WT) and LjNIN (RWP^LjNLP4^), though it retained substantial interaction with Pro*CLE-RS2*(i)/Pro*LjNIR1*(ii) (Fig. 1F). Conversely, LjNLP4 (FR^LjNIN^) bound strongly to Pro*LjNF-YA*(iii)/Pro*LjNF-YB*(iv) (Fig. 1F). A similar binding pattern of the chimeric proteins to *cis*-elements was also observed in other LjNIN target genes (fig. S3) (*5, 14, 17*). Chimeric proteins with both the RWP-RK and FR swapped (LjNIN (RWP^LjNLP4^ FR^LjNLP4^) /LjNLP4 (RWP^LjNIN^ FR^LjNIN^)) exhibited completely reversed DNA-binding specificities (Fig. 1G and fig. S2, C and D). Overall, these results indicate that the identified 15-residue FR^NIN^ is critical for the broad DNA-binding specificity of NIN.

We next used LjNIN/LjNLP4 deletion constructs lacking the C-terminal PB1 domain to assess the biochemical role of FR^NIN^ within a minimal DNA-binding module comprising only the RWP-RK and FR. EMSA showed that these modules, including FR-swapped chimeras, retained DNA-binding specificity nearly identical to the PB1-containing proteins, each producing a single shifted band regardless of protein concentration (Fig. 2, A to D, fig. S4). These findings indicate that the DNA-binding modules are sufficient to recapitulate the DNA-binding properties of LjNIN and LjNLP4, demonstrating the broad DNA-binding specificity conferred by FR^NIN^ is functionally active even in this minimal context.

**Fig. 2.**
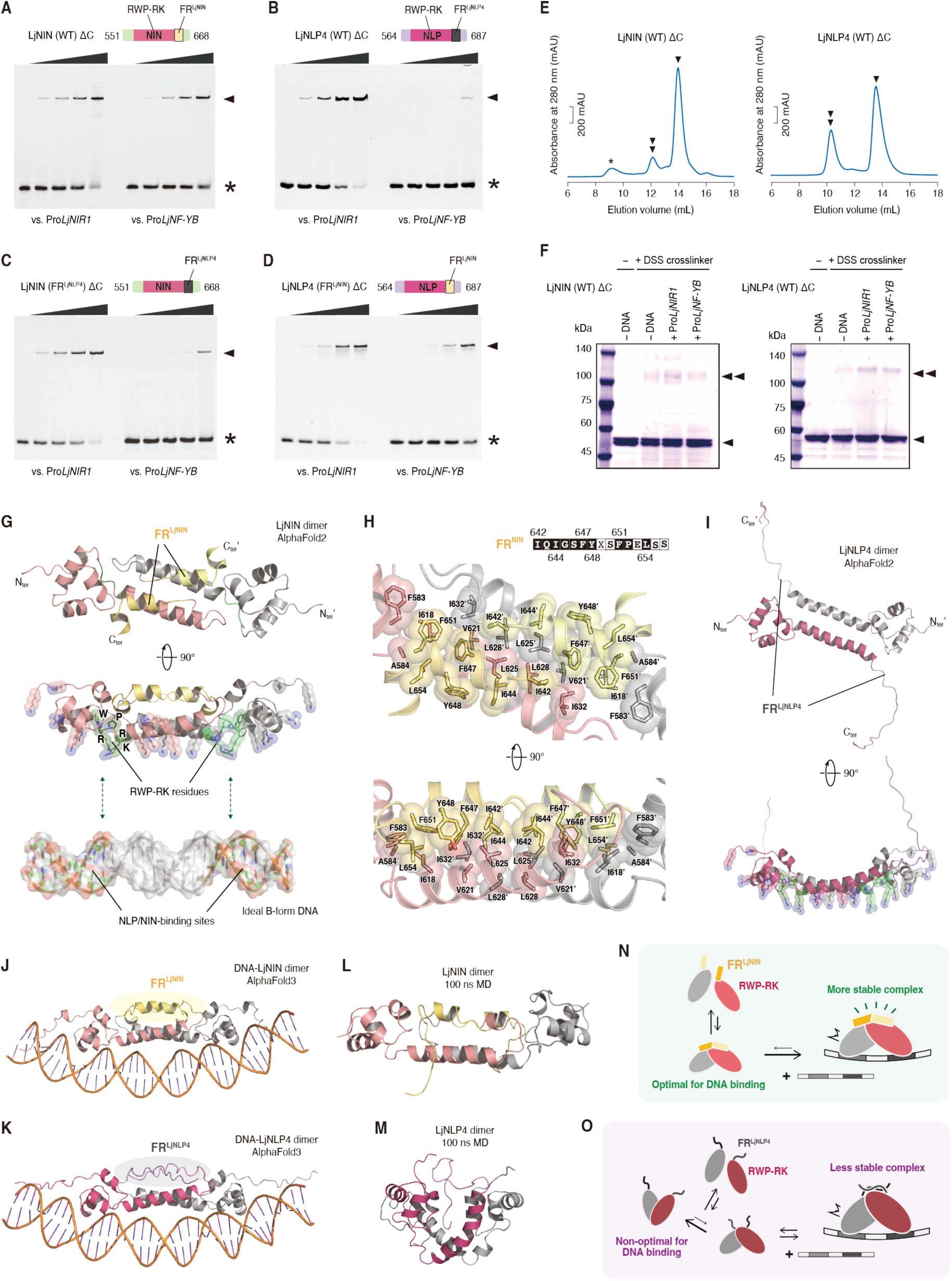
Molecular properties of the DNA-binding modules, including the RWP-RK and FR. (**A** to **D**) EMSA results of the LjNIN and NLP4 proteins with the C-terminus region deleted (LjNIN (WT) ΔC; A), LjNLP4 (WT) ΔC; B), and their FR-swapped chimeras (LjNIN (FR^LjNLP4^) ΔC; C), LjNLP4 (FR^LjNIN^) ΔC; D). The ΔC proteins fused to the MBP at N-terminus were reacted at 0.25, 0.5, 1 and 2 μM final conc. with 0.25 μM DNA probes. The electrophoretic patterns are shown with asterisks indicating the positions of free DNA. These experiments were repeated independently with similar results at least three times. (**E**) SEC analyses on the MBP-fused LjNIN (WT) ΔC and LjNLP4 (WT) ΔC. Chromatograms show monomeric and dimeric forms indicated by single and double arrowheads, respectively. Asterisk represents protein aggregation. (**F**) Chemical crosslink analyses of MBP-fused LjNIN (WT) ΔC and LjNLP4 (WT) ΔC using the amine-specific crosslinker DSS. SDS-PAGE results are shown for these proteins under DSS-treated and DSS-free conditions, as well as under DSS-treated conditions in the presence of target DNA including Pro*LjNIR1* and Pro*LjNF-YB*. The molecular masses of MBP-fused LjNIN (WT) ΔC and LjNLP4 (WT) ΔC are approximately 56 kDa and 57 kDa, respectively. (**G**) Top-ranked AlphaFold2-predicted structures of the LjNIN dimers containing the RWP-RK and FR. Two different chains are depicted with different colors. The five conserved “RWP-RK” residues are depicted with green color. The FR^LjNIN^ are highlighted with yellow-based colors. The structure of ideal B-form DNA fragment is shown on the same length scale as the LjNIN predicted structure. The two core-sites of NIN/NLP-binding motif are depicted with green color. (**H**) Close-up view of the top-ranked AlphaFold2-predicted structure of the LjNIN dimer. The residues forming hydrophobic and/or van der Waals interactions on FR^LjNIN^ are shown as stick and sphere models. (**I**) Top-ranked AlphaFold2-predicted structures the LjNLP4 dimers containing the RWP-RK and FR. (**J** and **K**) The top-ranked AlphaFold3-predicted structures of LjNIN (J) and LjNLP4 (K) dimers in complex with target DNA (Pro*LjCLE-RS2* fragment). The FR^LjNIN^ are highlighted with yellow-based colors. (**L** and **M**) Results of 100 ns MD simulations based on the AlphaFold2-predicted structure of LjNIN (L), and LjNLP4 (M) in the DNA-free state. (**N** and **O**) Proposed model underling the DNA-binding specificity differences between LjNIN (N) and LjNLP4 (O). While both RWP-RK domains can weakly dimerize, LjNLP4 tends to form suboptimal dimers, leading to unstable or inefficient DNA binding (O). In contrast, the FR^LjNIN^ stabilizes an optimal dimer interface of RWP-RK, enhancing binding efficiency and retention even to unpreferred nucleobases (N). This structural support enables broader DNA-binding specificity of LjNIN.

### The DNA-binding modules commonly possess an intrinsic ability to form dimers

Given that the target DNA probes contained two core-sites but yielded only a single shifted band in EMSA (Fig. 2, A to D, fig. S4), we hypothesized that the DNA-binding modules, composed of RWP-RK and FR, have an intrinsic potential to form dimers. To assess their oligomerization state, we performed size-exclusion chromatography (SEC) and Blue-Native-PAGE, which revealed that the modules of LjNIN and LjNLP4, and their chimeras exist mainly as monomers, but partially dimerize even without DNA (Fig. 2E, fig. S5). Crosslinking experiments using amine-specific crosslinker disuccinimidyl suberate (DSS) indicated that these dimerization as enhanced by their target DNA (Fig. 2F, fig. S6). Collectively, these findings suggest that the DNA-binding modules intrinsically dimerize, and that this dimerization is stabilized upon binding to *cis*-elements with two core-sites, potentially providing a basis for the different DNA-binding specificities.

### FR^NIN^ is predicted to reinforce the RWP-RK dimerization interface for broad DNA-binding specificity

Since biochemical approaches left the mechanism by which FR^NIN^ broadens DNA-binding specificity elusive, we next aimed to explore this question from a structural biology perspective. To gain structural insights, we predicted the dimeric structures of the DNA-binding modules of LjNIN and LjNLP4 using AlphaFold2. From a total of 100 structures generated by two advanced modes, we selected the one with the highest confidence value (fig. S7, A and B). Both top-ranked predicted structure of LjNIN and LjNLP4 exhibited dimer formation primarily facilitated by the RWP-RK. (Fig. 2, G to I). In these models, both RWP-RK domains dimerized primarily through a pair of leucine zipper-like helices (Fig. 2, G and H), stabilized by hydrophobic and van der Waals interactions via highly conserved residues in the NLP/NIN family (Fig. 1C, Fig. 2H, fig S1c). Meanwhile, the fully conserved hydrophobic residues within FR^LjNIN^ interacted with the dimerization helices of RWP-RK, forming a short helix that further supported the dimer interface (Fig. 2H). However, the FR^LjNLP4^ was not predicted to contribute to RWP-RK dimerization (Fig. 2I). This expanded dimer interface was observed in additional predicted structures of the LjNLP4-chimera with FR^LjNIN^, but not in those of the LjNIN-chimera with FR^LjNLP4^ (fig. S7, C and D).

Additionally, basic residues were concentrated on the same side of the predicted RWP-RK dimer structures (Fig. 2, G and I), a feature typical of DNA-binding proteins. Of the five completely conserved residues from which the term “RWP-RK” is derived, three basic residues clustered on the potential DNA-binding surface, while the remaining tryptophan and proline residues maintained characteristic structural features (Fig. 2, G and I). The arrangement of two “RWP-RK” residues aligns well with recognition of two core-sites approximately 10 bp apart in the target DNA (Fig. 1B and Fig. 2, G and I). Therefore, the predicted dimerization mode may also reflect the DNA-bound state. The FR regions were positioned opposite the DNA-binding surface (Fig. 2, G and I), consistent with our previous results that the FRs do not alter nucleobase-recognition preferences (*14*) (Fig. 1B). Furthermore, these findings were corroborated by AlphaFold3 predictions of LjNIN/LjNLP4 dimers complexed with DNA, which were structurally reliable with well-supported geometry at both the dimer and protein–DNA interfaces, with the slight distortion observed in the bound DNA (Fig. 2, J and K, fig. S8). Collectively, these results suggest that the FR^LjNIN^ contributes an additional dimer interface between the RWP-RKs, stabilizing a dimeric state that is optimal for recognizing the *cis*-elements harboring two core-sites. To further investigate the DNA-free state, we conducted molecular dynamics (MD) simulations. LjNIN maintained a stable dimeric structure suitable for DNA binding (Fig. 2L, fig. S9A), whereas LjNLP4 lost this conformation despite remaining dimeric (Fig. 2M, fig. S9B). These findings suggest that FR^NIN^ reinforces the RWP-RK interface both before and after DNA engagement. Consistently, DSS crosslinking of DNA-bound dimeric forms of LjNLP4 partially enhanced binding to unpreferred *cis*-elements, possibly by mimicking the FR^NIN^-stabilized state (fig. S10).

Taken together, we propose a model explaining the specificity differences between LjNIN and LjNLP4: while both RWP-RK domains can weakly dimerize (Fig. 2, N and O), LjNLP4 tends to form non-optimal dimers, resulting in unstable or inefficient binding to the unpreferred *cis*-elements (Fig. 2O). In contrast, FR^LjNIN^ stabilizes an optimal dimer interface of RWP-RK, enhancing the DNA-binding efficiency and retention, even at unpreferred *cis*-elements (Fig. 2N). This structural reinforcement underlies the increased versatility and robustness of LjNIN-mediated DNA recognition.

### FR^NIN^ has an essential role in NIN to regulate RNS

To test the significance of FR^LjNIN^ in plants, we overexpressed the full-length chimeric proteins in *L. japonicus* hairy roots (fig. S11A). In line with the EMSA results, swapping FR^LjNIN^ and FR^LjNLP4^ resulted in a functional conversion of each protein in inducing the expression of *LjNF-YA* and *LjNF-YB* (fig. S11, B to E). Moreover, domain swapping including RWP-RK region exerted an even more pronounced effect. In addition, FR-deleted LjNIN (LjNIN (ΔFR)) largely lost the ability to bind to Pro*LjNF-YA*(iii)/Pro*LjNF-YB*(iv) and other LjNIN-specific *cis*-elements, while still interacting with Pro*CLE-RS2*(i)/Pro*LjNIR*1(ii), regardless of the presence or absence of the C-terminal PB1 domain (figs. S3 and S12, A to C). Furthermore, structural prediction combined with MD simulation showed that DNA-free LjNIN (ΔFR) transitioned into a conformation non-optimal for DNA binding (fig. S12D), like the case of LjNLP4 (Fig. 2M). Overexpression of full-length LjNIN (ΔFR) in *L. japonicus* did not induce the expression of *LjNF-YA* and *LjNF-YB* (fig. S13, A to C).

Next, we created an *L. japonicus* mutant (ΔFR *nin*) using the CRISPR-Cas9 system, which resulted in a 60-bp nucleotide deletion specifically removing the FR containing region without introducing a frameshift mutation (Fig. 3, A and B). EMSA revealed that the mutated NIN protein in ΔFR *nin* (LjNIN (ΔFR^CR^)) could not bind to NIN target sequences, similar to LjNIN (ΔFR) (figs. S12B). While most known *nin* mutants, such as *nin-9*, completely lack nodulation (*18*), the ΔFR *nin* mutants retained nodule formation but developed non-functional white nodules with no nitrogen-fixing activity (Fig. 3C to F and fig. S14A). Nodule sections from ΔFR *nin* showed that rhizobia were localized in specific areas rather than uniformly distributed as in WT nodules (Fig. 3, G and H). Transmission electron microscopy revealed that ΔFR *nin* nodule cells were deformed and had fewer rhizobia colonized compared to WT; some rhizobia were possibly present in intercellular spaces (Fig. 3, I to N). Additionally, epidermal infection threads (ITs) were rarely observed in ΔFR *nin* roots (Fig. 3, O to Q). Cortical IT formation was also severely impaired in ΔFR *nin*, but rhizobia were observed on the surface of nodule primordia (fig. S14, B and C). RT-qPCR analysis showed that, unlike in *nin-9* mutants, target genes of NIN, such as *LjNF-YA*, *LjNF-YB*, *LjEPR3*, and *LjRPG*, were induced in ΔFR *nin*, albeit with delayed and/or reduced expression compared to WT (Fig. 3R and fig. S14D). This delayed gene expression likely contributes to the delayed nodule development observed in ΔFR *nin*. In *M. truncatula*, NIN directly regulates *Leghemoglobin* (*Lb*) gene expression (*19*). Consistent with the formation of non-functional nodules, *LjLb2* was not expressed at any tested stage in ΔFR *nin* (fig. S14D). We performed transcriptome analysis using roots of WT, the ΔFR *nin*, and the *nin-9* at 0, 7, and 21 dai. Among the genes upregulated by rhizobia in WT, 1,165 and 1,343 genes were downregulated in the ΔFR *nin* and the *nin-9*, respectively, at 7 dai (fig. S15A, tables S1 to S3). Among these, 1,050 genes were commonly downregulated in both mutants, suggesting that most function of NIN was attenuated in ΔFR *nin*. At 21 dai, 1,979 genes in the ΔFR *nin* and 2,414 genes in the *nin-9* were downregulated (fig. S15B, tables S4 to S6). Expression patterns of NIN target genes were consistent with that detected in RT-qPCR analysis (fig. S15C). In *M. truncatula nin-16* mutant with non-functional nodule formation, typically show upregulation of senescence- and defense-related genes (*20*), such activation was not observed in the ΔFR *nin* (fig. S15D).

**Fig. 3.**
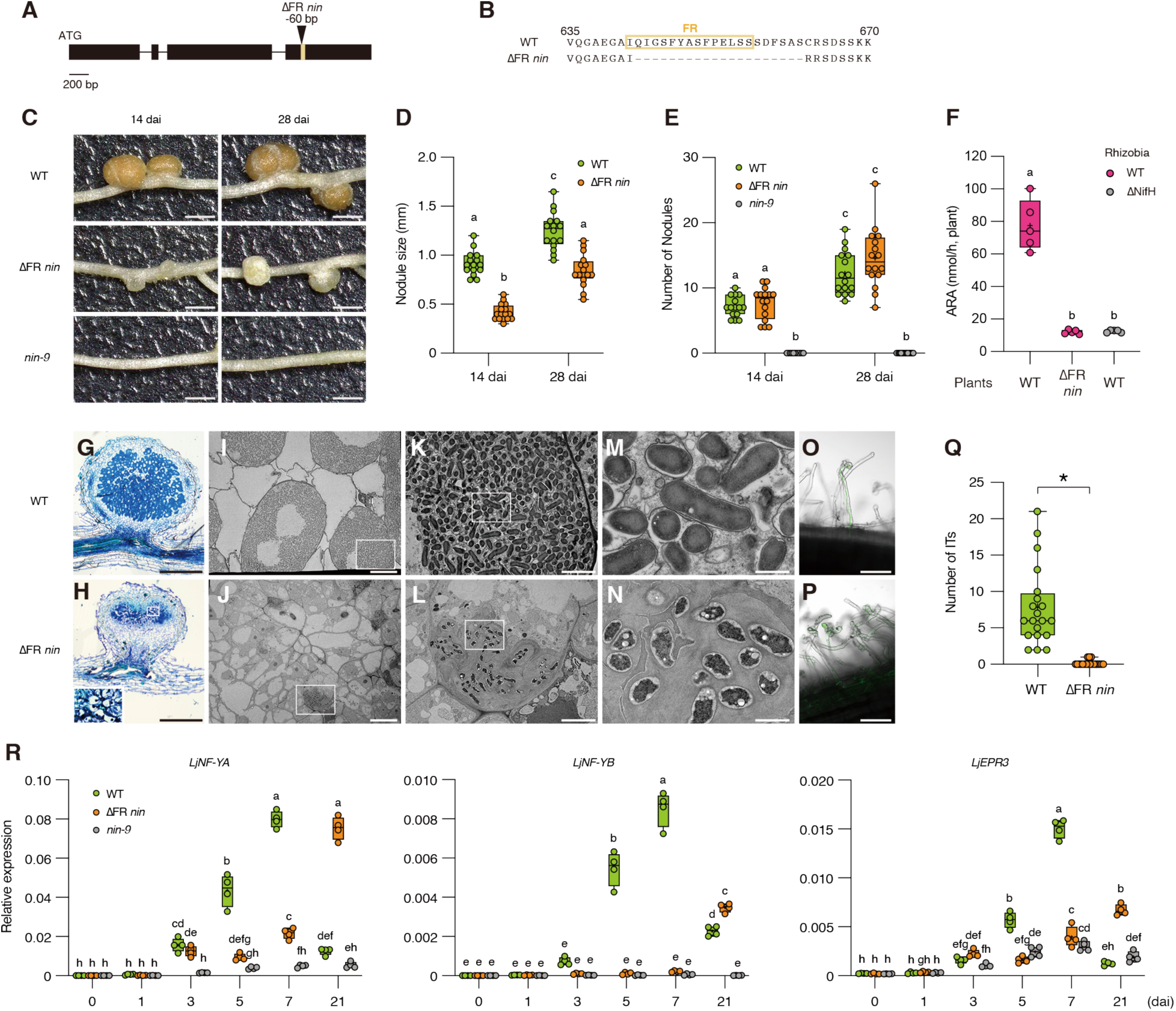
Phenotypes of ΔFR *nin* mutants. (**A**) Gene model of *LjNIN*. The black boxes and lines indicate exons and introns respectively. Arrowhead indicates the region that is deleted in ΔFR *nin* mutants. (**B**) Protein sequences of LjNIN around FR in WT and ΔFR *nin* mutants. (**C** to **E**) Nodulation phenotypes of ΔFR *nin* mutants at 14 and 28 days after inoculation (dai). (C) Nodules. (D) The diameter of the largest nodule in each plant. (E) Number of total nodules. (**F**) Acetylene reduction activity (ARA) of WT and ΔFR *nin* plants at 21 dai. WT plants inoculated with ΔNifH rhizobia (ΔNifH) were used as controls for the absence of nitrogen fixation. (**G** and **H**) Sections of nodules at 28 dai stained with toluidine blue. (G) WT, (H) ΔFR *nin*. The area indicated by the white square in (H) is enlarged to the left (**I** to **N**). Transmission electron microscopy images of nodule sections at 28 dai. The white squares in [(I), (J), (K), and (L)] indicate the area enlarged by [(K), (L), (M), and (N)]. [(I), (K), and (M)] WT, [(J), (L), and (N)] ΔFR *nin*. (**O** and **P**) Confocal images of representative infection threads (ITs) of WT at 7 dai (O) and ΔFR *nin* plants at 11 dai (P). (**Q**) Number of ITs formed in WT and ΔFR *nin* plants at 14 dai. (O to Q) Plants were inoculated with GFP-labelled rhizobia. (**R**) RT-qPCR analysis of *LjNF-YA*, *LjNF-YB*, and *LjEPR3* expressions in WT, ΔFR *nin*, and *nin-9* plants at 0, 1, 3, 5, 7, 21 dai. (n = 4, each n contains roots from at least 3 plants.) Data were normalized by *LjUBQ* expression. In box plots, individual biological replicates are shown as dots. In [(D), (E), and (R)], different letters indicate statistically significant differences (P < 0.05, Two-way ANOVA followed by multiple comparisons). In (F), different letters indicate statistically significant differences (P < 0.05, One-way ANOVA followed by multiple comparisons). In (Q), asterisk indicates a statistically significant difference (P < 0.05, Mann-Whitney test). Bars, 1 mm (C), 500 µm [(G) and (H)], 20 µm [(I) and (J)], 5 µm [(K) and (L)], 1 µm [(M) and (N)], and 100 µm [(O) and (P)].

We then examined spontaneous nodule formation by treating with the synthetic cytokinin 6-benzylaminopurine (BAP), or overexpressing a constitutively active form of CCaMK (T265D) (*21–23*). Spontaneous nodule formation was not induced in the ΔFR *nin* (fig. S16, A and B). Moreover, complementation analysis in the *daphne* mutant with defective in cortical NIN function (*24*), showed that LjNIN (ΔFR) induced fewer nodules and impaired nodule development compared to LjNIN (WT) (fig. S16C). These results suggest that the FR of NIN is required not only for rhizobial infection in the epidermis but also for the regulation of nodule organogenesis in the cortex. During the creation of ΔFR *nin*, we also obtained another *nin* mutant lacking the entire C-terminal region after RWP-RK, which completely abolished nodule formation (fig. S17).

### FR provides clues to the process by which NIN was derived from NLPs

Among the FRs of five LjNLPs, FR^LjNLP1^ appears to be most similar to FR^NIN^ (fig. S18A). To test functional equivalence of FR^LjNLPs^ to FR^LjNIN^, we performed complementation tests of the *nin-9* using chimeric proteins in which FR^LjNIN^ of LjNIN was swapped with each FR^LjNLPs^. Although ITs and immature nodules were occasionally induced by LjNIN with FRs from LjNLP2/3/4, only LjNIN (FR^LjNLP1^) formed mature pink nodules with shoots recovery comparable to that of LjNIN (FR^LjNIN^, WT) (fig. S18B). We then examined the functionally equivalence of full-length LjNLP1 and LjNIN. Overexpression of *LjNLP1* in *L. japonicus* hairy roots could induce the expression of *LjNF-YB*, but not *LjNF-YA* or *LjEPR3* (fig. S19, A and B). To avoid nitrate-induced suppression of nodule formation by LjNLP1/4 during activation of LjNLP1 by nitrate (*14, 25*), *nin-9 Ljnlp1 Ljnlp4-1* triple mutant was used in the complementation test. *LjNLP1* expressed under the *LjNIN* promoter fragment did not rescue the nodulation defects of *the triple* mutants (fig. S19C). Thus, these results suggest functional conservation of the FR alone is not sufficient for NLP to acquire NIN function. Consistent with the FR-swapped complementation tests, predicted structure of FR^LjNLP1^ was very similar to that of FR^LjNIN^ (fig. S20, A and B); however, the DNA-binding specificity of LjNLP1 differed from those of LjNIN and LjNLP4, exhibiting a distinct pattern of broad selectivity (Fig. 1D, fig. S20C). In contrast, the FR-swapped chimera LjNIN (FR^LjNLP1^) exhibited a DNA-binding specificity more similar to LjNIN (Fig. 1D, fig. S20D). These results suggest that the specificity of the FR-independent RWP-RK^LjNLP1^ is distinct from that of LjNIN, which may partly explain why LjNLP1 could not rescue the function of LjNIN.

To investigate the functional relevance of NIN-type FR in non-nodulating plants, we next examined AtNLP2, the evolutionarily closest homolog of LjNIN among the nine NLP family members in *Arabidopsis thaliana* (fig. S1, A and B). AtNLP2, in particular, exhibited broad DNA-binding specificity similar to LjNIN, whereas AtNLP2 lacking FR (AtNLP2 (ΔFR)) and AtNLP7, an orthologue of LjNLP4 (*25, 26*), specifically bound to perfect *cis*-elements (Fig. 4A). The FR-dependent expansion of the dimer interface was predicted in the dimeric structure of AtNLP2, but not in that of AtNLP7 (Fig. 4B and fig. S21A). In addition, the RWP-RK domain of AtNLP2 is more highly conserved with those of NINs than with other NLPs including LjNLP1 and LjNLP4, AtNLP7 (fig. S21B). These findings suggest that AtNLP2, unlike LjNLP1, shares biochemical features with LjNIN not only in the FR but also in the RWP-RK. Based on these observations, we hypothesized that AtNLP2 could mimic the biological function of LjNIN. Indeed, overexpression of *AtNLP2* in *L. japonicus* resulted in excessive root deformation, reminiscent of the effects of *LjNIN* overexpression (fig. S22A). Furthermore, in the absence of rhizobia, *AtNLP2* overexpression could induce the expression of LjNIN target genes, including *LjNF-YA*, *LjNF-YB*, and *LjEPR3* (Fig. 4, C and D). Consistent with the previous report, AtNLP2 activity was dependent on nitrate (*27*) (fig. S22, B and C). Of note, we found that *AtNLP2* expressed under *LjNIN* promoter could induce ITs and nodule formation in *nin-9 Ljnlp1 Ljnlp4-1* mutants in the presence of low nitrate in an FR dependent manner (Fig. 4E), although the nodules induced by AtNLP2 were imperfect in terms of accommodating rhizobia in nodules (Fig. 4F). Taken together, NLPs from non-nodulating plants that are evolutionarily close to NIN, such as AtNLP2, likely share fundamental functions with NIN. It may be primarily due to similar DNA-binding specificity conferred by the combination of RWP-RK and FR, though not entirely identical.

**Fig. 4.**
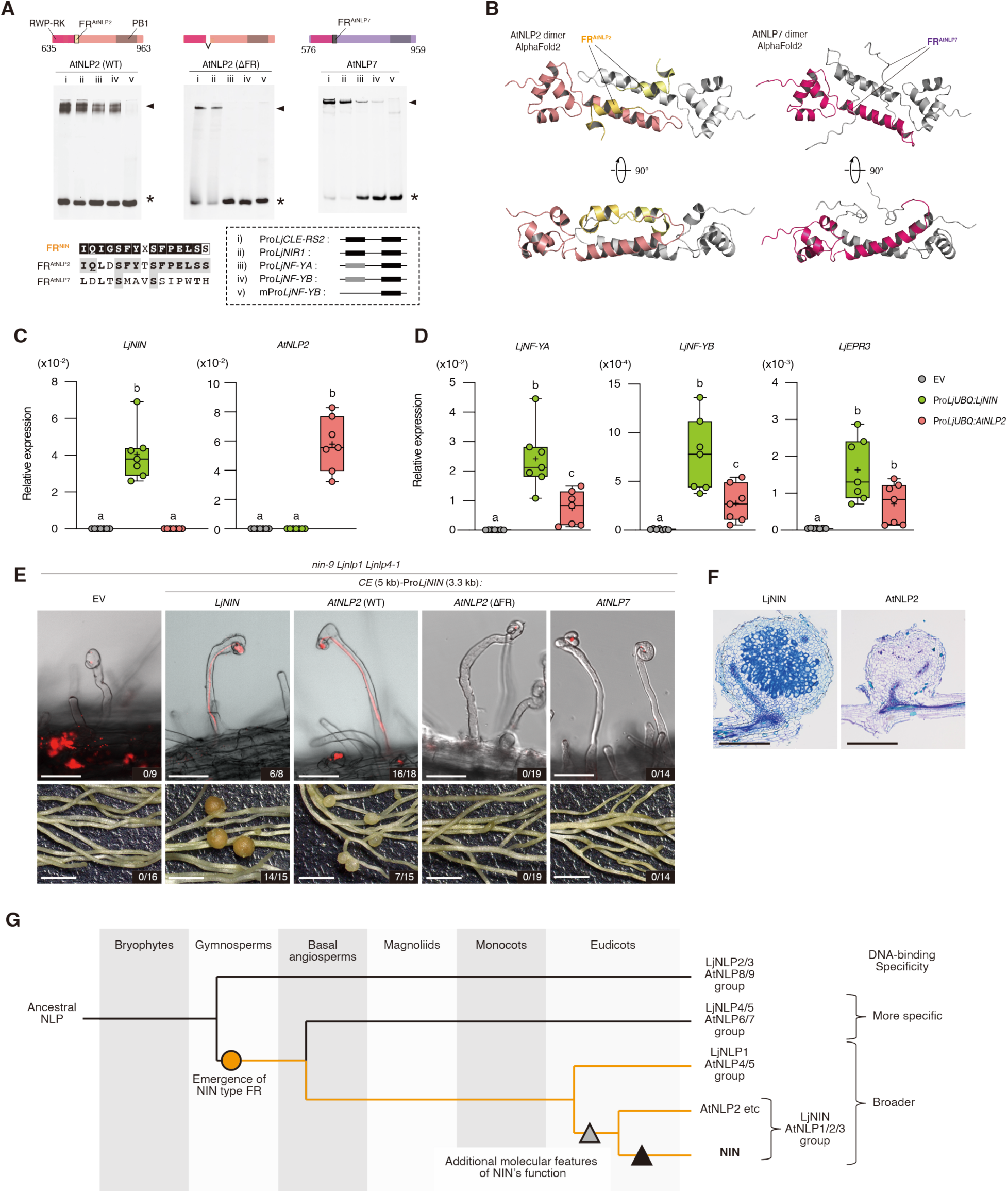
AtNLP2 shares fundamental features with NIN. (**A**) EMSA results of AtNLP2 with or without FR and AtNLP7. The alignment of FR and constructs of AtNLP2 and AtNLP7 proteins are shown at upper panels. Protein sequence alignment of FR^AtNLP2^ and FR^AtNLP7^ are shown. Highlighted letters on a gray background show the same amino acids as FR^NIN^. Boldfaces on white background in protein sequence alignment of FR^AtNLP2/7^ show amino acids with similar properties to those of FR^NIN^. Each recombinant protein fused to the MBP at N-terminus was reacted at 2 μM final conc. with 0.25 μM DNA probe (i to v; fig. S2A). The electrophoretic patterns are shown with asterisks and arrowheads indicating the positions of free DNA and the protein-DNA complexes, respectively. (**B**) Top-ranked AlphaFold2-predicted structures of the AtNLP2 and AtNLP7 dimers containing the RWP-RK and the FR. Two different chains are depicted with different colors. The FR^AtNLP2^ are highlighted with yellow-based colors. (**C** and **D**) RT-qPCR analysis of *LjNIN*, *AtNLP2*, *LjNF-YA*, *LjNF-YB*, and *LjEPR3* expressions in the transgenic hairy roots overexpressing *LjNIN* or *AtNLP2* by *LjUBQ* promoter. Transgenic plants were grown with 0.5 mM KNO_3_ in the absence of rhizobia for 1 week. EV: empty vector (n = 7, each n contains hairy roots from 3 transgenic plants). Data were normalized by *LjUBQ* expression. Individual biological replicates are shown as dots. Different letters indicate statistically significant differences (P < 0.05, One-way ANOVA followed by multiple comparisons). (**E** and **F**) *LjNIN*, *AtNLP2*, *AtNLP2* (*ΔFR*), or *AtNLP7* were expressed under *CE* (5 kb)-*LjNIN* promoter (3.3 kb) in hairy roots of *nin-9 Ljnlp1 Ljnlp4-1* triple mutants. Transgenic plants were inoculated with DsRED-labelled or WT rhizobia in the presence of 5 mM KNO_3_ for 30-38 d. (E) ITs (upper panels) and nodules (lower panels). Numbers indicate the frequency of plants forming ITs (upper panels) or nodules (lower panels) among all transgenic plants. (F) Sections of nodules induced by LjNIN or AtNLP2 were stained with toluidine blue. (**G**) **A model for the evolution of NIN and its DNA-binding specificity from NLP.** The NLP family underwent multiple duplications during land plant evolution, resulting in functional diversification (*11*, *33*, *34*)). The closest homologs of NIN in angiosperms are found in two groups: LjNIN and AtNLP1/2/3, and LjNLP1 and AtNLP4/5. These groups are absent in gymnosperms; however, *Cryptomeria japonica*, a gymnosperm species, possesses an NLP that includes a functional FR motif resembling the FR motif found in NIN (hereafter referred to as “NIN-type FR”). In contrast, MpNLP, an ancestral NLP from *Marchantia polymorpha*, a bryophyte species, lacks the NIN-type FR, suggesting that this motif is not an ancestral feature of the NLP family. Rather, it may have originated in the NLPs of gymnosperms. The dimeric conformation of the RWP-RK domain facilitated by the NIN-type FR confers broader DNA-binding specificity compared to NLPs lacking this motif. This structural feature may have played a key role in the functional divergence of NIN. The LjNIN and AtNLP1/2/3 group likely emerged in early eudicots, from which NIN was later derived within the nitrogen-fixing clade. Over the course of evolution, NIN appears to have acquired additional molecular features essential for its function, beyond just the NIN-type FR. Orange lines indicate NLPs containing the NIN-type FR. Bars, 50 µm (E, upper panels), 2 mm (E, lower panels), 500 µm (F).

Lastly, to gain insights into the origin of NIN-type FR, we collected NLPs that are potentially orthologous to NIN from eudicots, monocot, magnoliid, basal angiosperm, gymnosperm, and bryophyte. (fig. S23, A and B). Full length LjNIN with these FRs, except for FR^MpNLP^, induced ITs and functional mature pink nodule formation, although with varying efficiency (fig. S23C). Therefore, the acquisition of NIN-type FR might have occurred by the time gymnosperms emerged, during the molecular evolution of NLPs.

## Discussion

In this study, we identified FR^NIN^ as a crucial element responsible for NIN’s unique DNA-binding specificity. We also propose a fundamental mechanism of DNA recognition within the NLP family, driven by dual dimerization through multiple domains. The discovery of FR^NIN^ exemplifies how a TF can broaden its selectivity without altering its overall preference. To date, NLPs are known to act as dimers through the PB1 domain (*28*). Here, we demonstrated that the RWP-RK domains also weakly form dimers, where FR^NIN^ enhances NIN to bind to unpreferred *cis*-elements by sustaining the dimerization of the RWP-RK domains. Although the ΔFR *nin* severely impairs rhizobial infection and nodule organogenesis, some RNS processes, including nodule initiation, are still maintained, indicating NIN’s role is not completely attenuated in ΔFR *nin*. One possible explanation is that FR^NIN^ functions primarily to stabilize the NIN-dimer binding to DNA in a context dependent manner, which may change spatiotemporally following nodule development. Under conditions in which FR^NIN^ is dysfunctional, there appears to be a mechanism by which NIN can form a dimer through other domains and regulates the expression of its target genes. In particular, considering the expression of *LjNF-YA*, NIN may have a function that does not require FR^NIN^. Furthermore, when NIN function is attenuated, other factors may regulate the expression of NIN target genes through a NIN-independent mechanism.

Previous studies have shown that several prerequisites contributed to the establishment of NIN as the key TF regulating RNS. These include the evolution of the *NIN* promoter, which enabled its regulation during RNS, and the emergence of NIN-binding site in the promoters of many RNS-related genes (*4*, *29*-*31*). In this study, we identified an additional prerequisite related to NIN protein function. NIN-type FR, the motif responsible for its broad DNA-binding capacity, which is one of NIN’s defining features, had already appeared in certain NLPs before the evolution of RNS. It is likely that NLPs with functional FR similar to FR^NIN^ had already emerged at least as early as in gymnosperms (Fig. 4I). Elucidating the function of NIN-type FR in those NLPs remains an intriguing subject for future research. While intact AtNLP2 can substitute for many functions of LjNIN, it does not fully replicate LjNIN’s role. This finding suggests that additional molecular characteristics, such as the loss of nitrate responsiveness and/or other yet unidentified features, are likely critical for the evolutionary transition from NLPs to a fully functional NIN.

## Acknowledgments

We thank Gene Research Center, University of Tsukuba for technical support of the gel imaging, and M. Ohtsuka and K. Miura (University of Tsukuba) for technical support of the protein preparation.

## Funding

Ministry of Education, Culture, Sports, Science and Technology KAKENHI grant JP22K14824 and JP24K01677 (S.N.); JP20H05908, JP23K27188 and JP25H01345 (T.S.)

JST Mirai Program JPMJMI20E4 (T.S.) JST ALCA-Next JPMJAN23D2 (T.S.)

JST ACT-X Program JPMJAX21BL (S.N.) JST SPRING JPMJSP2124 (M.N.)

## Author contributions

Conceptualization: S.N., T.S.

Data curation: S.N., M.N., M.I.

Formal analysis: S.N., M.N., H.O., M.I.

Funding acquisition: S.N., M.N., T.S.

Investigation: S.N., M.N., H.O., M.I., T.S.

Methodology: S.N., M.N., T.S.

Project administration: S.N., T.S.

Resources: S.N., T.S.

Supervision: T.S.

Validation: S.N., M.N., T.S.

Visualization: M.N.

Writing – original draft: S.N., M.N., T.S.

Writing – review & editing: S.N., M.N., H.O., M.I., T.S.

## Notes

### Competing Interest Statement

The authors have declared no competing interest.

