## Supplementary Materials for "An NLP-inherited motif confers broad DNA-binding specificity to NIN in root nodule symbiosis"

**The PDF file includes:**

Materials and Methods
Figs. S1 to S23
Tables S1 to S7
References

### Materials and Methods

Phylogenetic analyses

Phylogenetic analyses in fig. S1, A and B were conducted in Mega X (*35*, *36*) using amino acid sequences of the full-length and DNA-binding regions from the 49 and 48 NIN/NLP family proteins, respectively. The evolutionary history was inferred using the Neighbor-Joining method (*37*). The optimal trees with the sum of branch length = 12.22349058 and 7.82173710 are shown for the full-length and DNA-binding regions, respectively. The percentage of replicate trees in which the associated taxa clustered together in the bootstrap test (1000 replicates) is shown next to the branches (*38*). The trees are drawn to scale, with branch lengths in the same units as those of the evolutionary distances used to infer the phylogenetic trees. The evolutionary distances were computed using the Poisson correction method (*39*) and are in the units of the number of amino acid substitutions per site. All ambiguous positions were removed for each sequence pair (pairwise deletion option). There were a total of 1531 and 156 positions in the final dataset the full-length and DNA-binding regions, respectively. Phylogenetic analysis in fig. S23A was conducted in MEGA11 (*40*) using amino acid sequences of DNA-binding regions from the 29 NIN/NLP family proteins. The tree was constructed by maximum likelihood (ML) method using IQ-TREE 2 (*41*) (JTT+F+I+G4 substitution model). The optimal tree with the sum of branch length = 25.915. The percentage of replicate trees in which the associated taxa clustered together in the bootstrap test (1000 replicates) is shown next to the branches. The tree was rooted to the outgroup MpNLP and drawn to scale, with branch lengths in the same units as those of the evolutionary distances used to infer the phylogenetic trees. The evolutionary distances were computed using the Poisson correction method and were in the units of the number of amino acid substitutions per site.

Sequence alignments

CLUSTAL OMEGA (*42*) was used for multiple sequence alignments among NIN/NLP TFs using default parameters, and the results were displayed by ESPript 3.0 (*43*). Aligned protein sequences included PaNIN from *P. andersonii* (accession: PON66248.1), and CgNIN from *C. glauca* (accession: KF481969.1), in addition to those used in the phylogenetic analyses (fig. S1).

Protein expression and purification

MBP-fused recombinant proteins for EMSA, SEC, Blue Native PAGE, and crosslinking experiments were prepared based on the method previously described (*14*) with some modifications. All truncated and chimeric constructs were BamHI-introduced into the pMAL-c2X vector (New England Biolabs) and cloned to Rosetta 2 (DE3) (Novagen). Overexpression was induced by 0.25 mM isopropyl-β-d-(-)-thiogalactopyranoside (IPTG) at 18 °C for 16 hr. For constructs containing the C-terminal region, cultures were continuously shaken at 110 rpm throughout the 16-hr induction at 18 °C. In contrast, for ΔC constructs, cultures were kept stationary for 12 hr and then shaken at 110 rpm for an additional 4 hr to improve the yield of soluble and active protein. After incubation, cells were harvested by centrifugation at 2750 ×g for 15 min and stored at −80 °C until use. Harvested cells were resuspended in Binding buffer. For constructs containing the C-terminal region, the buffer contained 20 mM Tris-HCl, pH 8.0, 1.0 M NaCl, 1 mM DTT and 10% glycerol, 0.5 mM EDTA with the addition of 0.1% (v/v) protease inhibitor cocktail (Nacalai Tesque). For ΔC constructs, the buffer was identical except that DTT was omitted and 1 mM PMSF was added. The resuspended cells were stirred with 0.02% (w/v) lysozyme and then lysed by sonication. The resuspended cells were stirred with 0.02% (w/v) lysozyme, and then lysed by sonication. The cell debris was removed thoroughly by centrifugation at 40,000 ×g for 30 min at 4 °C. Crude protein fractions were applied to fresh (non-regenerated) Amylose Resin (New England Biolabs), washed with the Binding buffer, and then eluted with Elution buffer (20 mM Tris-HCl, pH 8.0, 1.0 M NaCl, 1 mM DTT, 10% glycerol, 0.5 mM EDTA and 30 mM maltose) with the addition of 0.1% (v/v) protease inhibitor cocktail to obtain MBP-tagged proteins. All eluted fractions were concentrated by VIVASPIN Turbo (30,000 MWCO PES) (Sartorius) and further purified by size exclusion chromatography (SEC) using a Superdex 200 10/300 GL increase column (Cytiva) in an ÄKTA go protein purification system (Cytiva). SEC was performed using SEC buffer (20 mM Tris-HCl, pH 8.0, 1.0 M NaCl, 1 mM DTT, and 10% glycerol) at 0.4 ml/min flow rate. Purified and unaggregated proteins were re-concentrated and measured for concentration by their absorbance at 280 nm using NanoDrop One (Thermo Fisher Scientific) using their corresponding molecular weight and molar extinction coefficient at 280 nm.

EMSA

Interaction analyses of the NIN/NLP family proteins-DNA were performed based on the method previously described (*14, 44*) with some modifications. To prepare the probes, DNA fragments were labeled with carboxyfluorescein (FAM). The labeled DNA fragments were annealed in TNE buffer (20 mM Tris-HCl, pH 8.0, 200 mM NaCl, and 1 mM EDTA), and then purified on the Superdex 200 increase 10/300 GL in the ÄKTA go. In a total of 20 µl, each purified DNA fragment (0.25 µM) and poly(dI-dC) (50 ng/µL) were mixed with the purified proteins in EMSA reaction buffer (10 mM Tris–HCl pH 7.5, 50 mM KCl, 50 mM NaCl, 1 mM DTT, 2.5% glycerol, and 5 mM $MgCl_2$), and incubated at 25 °C for 20 min. 10 µl of each mixture was loaded on a 10% polyacrylamide gel in 0.5×TBE buffer (45 mM Tris–HCl, pH 8.2, 45 mM boric acid, and 1 mM EDTA). Fluorescence intensity (FI) was detected after 1 s exposure with the preset settings (Epi-blue excitation 466 nm, BPF535 bandpass filter 535 nm) using LuminoGraph III WSE-6300 (ATTO). The complex ratio of the protein–DNA complex (%) (= 100 × FIcomplex / (FIfree-DNA + FIcomplex)) was calculated using the Image J (Ver. 1.53a) application.

SEC for oligomeric analysis

For oligomeric analysis, MBP-fused ΔC proteins were preincubated at a final concentration of 500 µM on ice for one week. Immediately prior to SEC, each sample was diluted to 60 µM in a total volume of 250 µl, SEC was performed using the Superdex 200 Increase 10/300 GL column at 4 °C, with a flow rate of 0.4 mL/min. The SEC buffer consisted of 20 mM Tris-HCl pH 8.0, 1.0 M NaCl, 1 mM DTT, and 10% glycerol. The elution was monitored at 280 nm using the ÄKTA go system.

Blue Native PAGE

Purified proteins were mixed with an EzApply Native (ATTO), and then separated on a u-PAGEL H 4-20% gradient polyacrylamide gel (ATTO) in an EzRun BlueNative buffer (ATTO). The CBB-stained gel was destained with microwave heating and photographed. For comparison, the same protein samples were separated on an e-PAGEL 3-14% gradient polyacrylamide gel (ATTO) by SDS-PAGE and stained with CBB.

Crosslinking experiments

Prior to crosslinking with amine-specific crosslinker disuccinimidyl suberate (DSS), MBP-fused ΔC proteins, and MBP alone were buffer-exchanged into 20 mM HEPES-NaOH pH 7.5, 1.0 M

NaCl, 1 mM DTT, and 10% glycerol using the Superdex 200 Increase 10/300 GL column, in order to remove Tris buffer, which interferes with the DSS reaction. For SDS-PAGE-based crosslinking analysis, 10 µL reactions were prepared containing either 2 µM prepared protein with 1 µM non-labeled DNA fragment, or protein alone (no DNA condition), in a buffer composed of 10 mM HEPES-NaOH pH 7.5, 100 mM NaCl, 1 mM DTT, 2.5% glycerol, and 5 mM $MgCl_2$. After incubation at 25 °C for 10 min, disuccinimidyl suberate (DSS) was added to a final concentration of 200 µM (2% DMSO), and the reaction was continued at 25 °C for 20 min. Reactions were quenched by adding Tris-HCl (pH 8.0) to a final concentration of 50 mM. The entire volume of each reaction sample was loaded onto the e-PAGEL 3–14% gradient polyacrylamide gel for SDS-PAGE. The gel was stained with 0.25% CBB R-250 in 45% methanol and 10% acetic acid for 5 minutes, then replaced with water and heated in a microwave for 10 minutes, followed by destaining in water for two days. The ratio of DSS-mediated crosslinked dimers (%) was calculated as follows: 100 × (intensity of the crosslinked dimer band) / (intensity of the crosslinked dimer band + intensity of the non-crosslinked band), based on densitometric analysis of CBB-stained SDS-PAGE gels using the ImageJ (version 1.53a). For EMSA-based crosslinking analysis, 10 µl reactions were prepared containing 0, 1, or 2 µM protein with 0.25 µM FAM-labeled DNA fragment and 50 ng/µL poly(dI-dC), in the same buffer as above. After incubation at 25 °C for 10 min, DSS was added to a final concentration of 200 µM (2% DMSO), and reactions were continued at 25 °C for 20 min. For DMSO controls, 2% DMSO was added instead of DSS. Reactions were quenched by adding Tris-HCl (pH 8.0) to a final concentration of 50 mM, and the entire sample was analyzed under the same conditions described in the EMSA section.

Structure predictions

The protein dimer structures of the DNA-binding modules (including both RWP-RK and FR) of LjNIN, LjNLP4, LjNLP1, AtNLP2, and AtNLP7 were predicted under two conditions using ColabFold v.1.0.0 with the AlphaFold v.2.1.0 monomer-ptm model (ptm) or LocalColabFold v.1.5.2 with the AlphaFold v.2.3.0 'multimer_v3' model (mv3) (*45-47*). A total of 100 protein structures were predicted using 10 seed parameters, with 5 models generated for each condition. The average per-residue pLDDT score of the predicted full-length proteins indicated that the monomer-ptm model produced high-confidence predictions (Fig. S7A, B). The protein structures predicted by the monomer-ptm models were used for subsequent molecular dynamics (MD) simulations. These 100 protein structures were classified through principal component analysis based on the xyz-coordinates of all Cα atoms. AlphaFold3 predictions were performed by the AlphaFold server (https://alphafoldserver.com.) (*48*). For prediction, 2 copies of each DNA-binding module and Pro*LjCLE-RS2*-derived double-stranded DNA fragment (Fig. S2A) were simultaneously input. The predicted structures with pLDDT scores, and separate figures showing the corresponding PAE values were shown as the output from AlphaFold program, and those of the top ranked prediction without pLDDT were visualized in PyMOL 2.5.2 (Schrodinger, LLC) (*49*).

Molecular dynamics simulation

MD simulations were performed using the Maestro interface with the Desmond/GPU program (*50*). The three predicted protein structures were used as the initial templates for the MD simulations. Initial model of LjNIN with FR deleted was prepared by truncating FR and post-FR region from LjNIN models. The protonation states and orientations of the residues were determined by the Protein Preparation Wizard in Maestro using PROPKA (*51*). The solvent box was prepared in a cubic shape, with a TIP3P water model positioned 10 Å away from the protein surface (*52*). A 150 mM sodium chloride solution was added to the solvent box using the System Builder in Desmond (*53*). The force field for the protein atoms was assigned to OPLS-AA (all-atom) (*54*). After a 100 ps energy minimization step, a 100 ns MD simulation was performed under the following conditions: equilibration was conducted using the isothermal-isobaric ensemble (NPT). Short-range electrostatic interactions between atoms over 9 Å apart were cutoff, and long-range electrostatic interactions were computed using the Particle Mesh Ewald method. All covalent bonds were constrained using the SHAKE algorithm (*55*). The thermostat and barostat were Nose-Hoover chain (*56*) at 300 K and Martyna-Tobias-Klein (*57*) at 1 atm, with relaxation times of 1 ps and 2 ps, respectively. The RESPA integrator (*58*) was used, with Fourier-space electrostatics computed every 6 fs and all remaining interactions computed every 2 fs. Intermediate structures were recorded every 10 ps, and the Cα RMSD was calculated and analyzed relative to the initial structure.

Plant materials and growth conditions

The Miyakojima MG-20 ecotype of *L. japonicus* was used as the WT plant (*59*). A description of *nin-9*, *nlp1* and *nlp4-1* plants was published previously (*14, 18, 25*). Plants were grown with or without *Mesorhizobium loti* MAFF 303099 in autoclaved vermiculite or on 1% agar plates with Broughton and Dilworth (B&D) solution (*60*) under a 16 h light/8 h dark cycle at 24°C. ΔNifH strain of *M. loti* was previously reported (*61*). For the observation of rhizobial infection, plants were inoculated with *M. loti* MAFF 303099 constitutively expressing *GFP* or *DsRED*.

Stable and hairy root transformation

To create ΔFR *nin* plants by CRISPR–Cas9 system, gRNAs were designed using the CRISPR-P 2.0 program (fig. S17A and table S7) (*62*). Two gRNAs were used to target a gene, and they were cloned to pMR203_AB and pMR203_BC, gRNA cloning vectors, and then to pMR285_AD, a binary vector (*63*) by a standard ligation method using a DNA Ligation Kit (Takara) and several restriction enzymes. For stable transformation of *L. japonicus*, seeds were germinated on germination medium (1/2× Gamborg's B5 salt mixture (Wako), 1/2× Gamborg's vitamin solution (Sigma), 1% sucrose, 1% agar) in a growth cabinet (24 °C dark for first 2 d, 24 °C 16 h light/8 h dark cycle for next 2 d). *Agrobacterium tumefaciens* GV3101 MP90RK strains harboring a construct were streaked on YEP plate with appropriate antibiotics for 2 d at 28 °C. Seedlings were placed in the *A. tumefaciens* suspension, and then their hypocotyls were cut into about 3 mm pieces. The hypocotyl pieces were placed onto the top of pilled filter papers saturated with co-cultivation medium (1/10× Gamborg's B5 salt mixture, 1/10× Gamborg's vitamin solution, 0.2 μg $ml^{-1}$ BAP, 0.05 μg $ml^{-1}$ NAA, 5 mM MES (pH 5.2), 20 μg $ml^{-1}$ acetosyringone, pH 5.5) and were incubated in a growth cabinet (21 °C dark) for 6 d. After that, the hypocotyl pieces were transferred to callus induction medium (1× Gamborg's B5 salt mixture, 1× Gamborg's vitamin solution, 2% sucrose, 0.2 μg $ml^{-1}$ BAP, 0.05 μg $ml^{-1}$ NAA, 10 mM $(NH_4)_2SO_4$, 0.3% phytagel, 12.5 μg $ml^{-1}$ meropen, 15 μg $ml^{-1}$ Hygromycin B, pH 5.5) and were incubated in a growth cabinet (24 °C 16 h light/8 h dark cycle) for 2–3 w. The hypocotyl pieces were transferred to a fresh callus induction medium every 1 w. When calluses became more than 1 mm in size, they were detached from the hypocotyls and transferred onto callus medium without hygromycin B, and were incubated for 3–7 w in a growth cabinet (24 °C 16 h light/8 h dark cycle) until leaf primordia became visible. The calluses were transferred onto a new medium every 1 w. The calluses with leaf primordia then were transferred to shoot elongation medium (1× Gamborg's B5 salt mixture, 1× Gamborg's vitamin solution, 2% sucrose, 0.2 μg ml$^{-1}$ BAP, 0.3% phytagel, 12.5 μg ml$^{-1}$ meropen, pH 5.5), and incubated until their shoot length became about 1 cm. Individual shoots were detached from calluses and transferred to root induction medium (1/2× Gamborg's B5 salt mixture, 1/2× Gamborg's vitamin solution, 1% sucrose, 0.5 μg ml$^{-1}$ NAA, 0.4% phytagel, 12.5 μg ml$^{-1}$ meropen, pH 5.5), and incubated for 8 d. Then, they were transferred to a root induction medium without NAA and cultivated until their root length became about 2–3 cm. The resultant transgenic plants were transplanted into soils for further cultivation. Transgenic plants with homozygous mutations were used for analysis.

For *LjNIN*, *LjNLP4*, *LjNLP1*, or *AtNLP2* overexpression in *L. japonicus* hairy roots, the Pro*LjUBQ*-t*NOS* cassette was amplified by PCR from an original vector (*14*) and was cloned to pCAMBIA1300-GFP by the In-Fusion (Clontech) reaction. The coding sequence (CDS) of *LjNIN*, *LjNLP4*, *LjNLP1*, or *AtNLP2* was amplified by PCR and was cloned downstream of Pro*LjUBQ* by the In-Fusion reaction. Constructs described in Protein expression and purification were used as templates to amplify fragments of CDS of the RWP-RK and/or the FR swapped LjNIN/LjNLP4 proteins. For complementation of *nin* mutants, 5 kb region around *CE* (*30*) and Pro*LjNIN* 3.3 kb region upstream of start codon (*CE* (5 kb)-Pro*LjNIN* (3.3 kb)) (*31*), and tLjNIN were amplified by PCR from WT genomic DNA and were cloned to pCAMBIA1300-GFP by the In-Fusion reaction. Then, the coding sequence (CDS) of *LjNIN*, *LjNLP1*, *AtNLP2, or AtNLP7* was amplified by PCR and was cloned downstream of *CE* (5 kb)-Pro*LjNIN* (3.3 kb) by the In-Fusion reaction. To generate LjNIN (FR$^{swap}$) and AtNLP2 (ΔFR) constructs, two fragments of CDS of LjNIN or AtNLP2 before and after the FR were amplified by PCR using primers shown in table S7 and cloned downstream of *CE* (5 kb)-Pro*LjNIN* (3.3 kb) by the In-Fusion reaction. For hairy root transformation, seeds were germinated on the germination medium described above in a growth cabinet (24 °C dark for first 2 d, 24 °C 16 h light/8 h dark cycle for next 2 d). *A. rhizogenes* AR1193 strains harboring each construct were streaked on YEP plate with appropriate antibiotics for 2 d at 28 °C. Seedlings were placed in the *A. rhizogenes* suspension and then cut at the base of the hypocotyls. The seedlings with cotyledons were transferred onto hairy root medium (1× Gamborg's B5 salt mixture, 1× Gamborg's vitamin solution, 2% sucrose, 1% agar) and were grown in a growth cabinet (24 °C dark for first 1 d, 24 °C 16 h light/8 h dark cycle for next 2 d). Then, the plants were transferred onto fresh hairy root medium containing 12.5 μg ml$^{-1}$ meropen and were grown for 7-9 d in a growth cabinet (24 °C 16 h light/8 h dark cycle) (*64*). Transgenic roots were identified by GFP fluorescence, and the plants with transgenic hairy roots were used for further experiments.

Acetylene reduction assay

The nitrogenase activity of nodules was indirectly determined by measuring the acetylene reductase activity (*25*). Nodulated plants were put into 20 mL vials. Subsequently, acetylene was injected into the vials. After incubation for 10 min, the amount of ethylene produced was measured using a GC-2014 (Shimadzu).

Toluidine blue staining

Root segments with nodules were fixed at 4°C overnight with 4% paraformaldehyde, 0.25% glutaraldehyde in 0.05 M phosphate buffer (pH 7.2). The fixed material was dehydrated in an ethanol series and subsequently embedded in Technovit 7100 (Kulzer) according to the manufacturer's protocol. Longitudinal sections (8 µm) were made by using a RM2265 microtome (Leica Microsystems) and stained for 5 min in 0.05% Toluidine Blue O. Sections were analyzed by using a BX53 microscope equipped with a DP74 camera (OLYMPUS).

Transmission electron microscopy analysis

Nodules were cut from roots and immediately placed in a solution of 2% paraformaldehyde, 2% (v/v) glutaraldehyde in 0.05 M cacodylate buffer, pH 7.4 for fixation, and left overnight at 4°C. The fixative was washed out by three successive 30-min washes in 0.05 M cacodylate buffer and post- fixed in 2% OsO4 in 0.05 M cacodylate buffer at 4°C overnight. The fixed material was dehydrated in an ethanol series (50% and 70% each for 30 min at 4°C, 90% and three changes of 100% ethanol each for 30 min at rt, and 100% ethanol for 2 d at rt). The dehydrated samples were washed out by two successive 30- min propylene oxide (PO). The samples were infiltrated with Quetol-651 resin (Nisshin EM) by successive changes of resin:PO mixes at rt (1:1 for 3h and 100% resin for 3h) then 100% resin at 60°C for 48 h to polymerize. The material was sectioned with a diamond knife using a Leica Ultracut UCT ultramicrotome (Leica) and ultrathin sections of approximately 80 nm were transferred onto copper grids. The sections were stained with 2% uranyl acetate for 15 min at rt and Lead stain solution (Sigma-Aldrich) for 3 min. The grids were viewed in a JEM-1400Plus (JEOL) at 100 kV and imaged using a CCD camera EM-14830RUBY2 (JEOL).

Gene expression analysis

The primers used for PCR are listed in Table S1. Total RNA was isolated from respective tissues using the PureLink Plant RNA Reagent (Invitrogen) or the Plant Total RNA Mini Kit (Favorgen Biotech). First-strand cDNA was prepared using the ReverTra Ace qPCR RT Master Mix with gDNA Remover (Toyobo). RT-qPCR was performed using a CFX Opus 384 Real-Time PCR system (BIO-RAD) with a THUNDERBIRD SYBR qPCR Mix (Toyobo) or a THUNDERBIRD Next SYBR qPCR Mix (Toyobo) following the manufacturer's instructions.

RNA-seq analysis

Total RNA was isolated from roots using the PureLink Plant RNA Reagent (Invitrogen). Libraries were prepared using a NEBNext Ultra II RNA Library Prep Kit from Illumina (New England Biolabs) following the manufacturer's instructions and sequenced using a Novaseq 6000 (Illumina) instrument with the 150-bp paired-end sequencing protocol. RNA-seq reads were mapped to the L. japonicus MG-20 genome version 3.0 using HISAT2 (ver. 2.2.1) with the default parameters (*65*). Mapped reads were then assembled using StringTie (ver.1.3.5) (*66*). Up- or down-regulated genes were identified with edgeR (ver. 3.40.2) (*67*), using a 5% false discovery rate (FDR). Only genes with a count per million (CPM) >= 0.5 in at least three out of the six samples derived from two different conditions were included in each analysis.

Microscopy

Bright-field images were taken under a S9i stereo microscope (Leica) or a BX53 upright microscope (OLYMPUS). Fluorescent images were obtained using a LSM700 confocal laser-scanning microscope (Carl Zeiss) equipped with ZEN (Carl Zeiss), a M205FA fluorescence stereo microscope (Leica), or a DM6 B upright microscope (Leica) equipped with LAS X software which processes images following THUNDER algorisms (Leica).

Statistical analysis

Statistical analysis was performed using GraphPad Prism version 9 (GraphPad Software). Normality was checked using the Shapiro–Wilk test and $P > 0.05$ was considered as normal distribution. The F-test was used to test whether the variances of the two populations were equal.

Appropriate methods were chosen according to the nature of the data. The criterion of $P < 0.05$ means a statistically significant difference in this study.

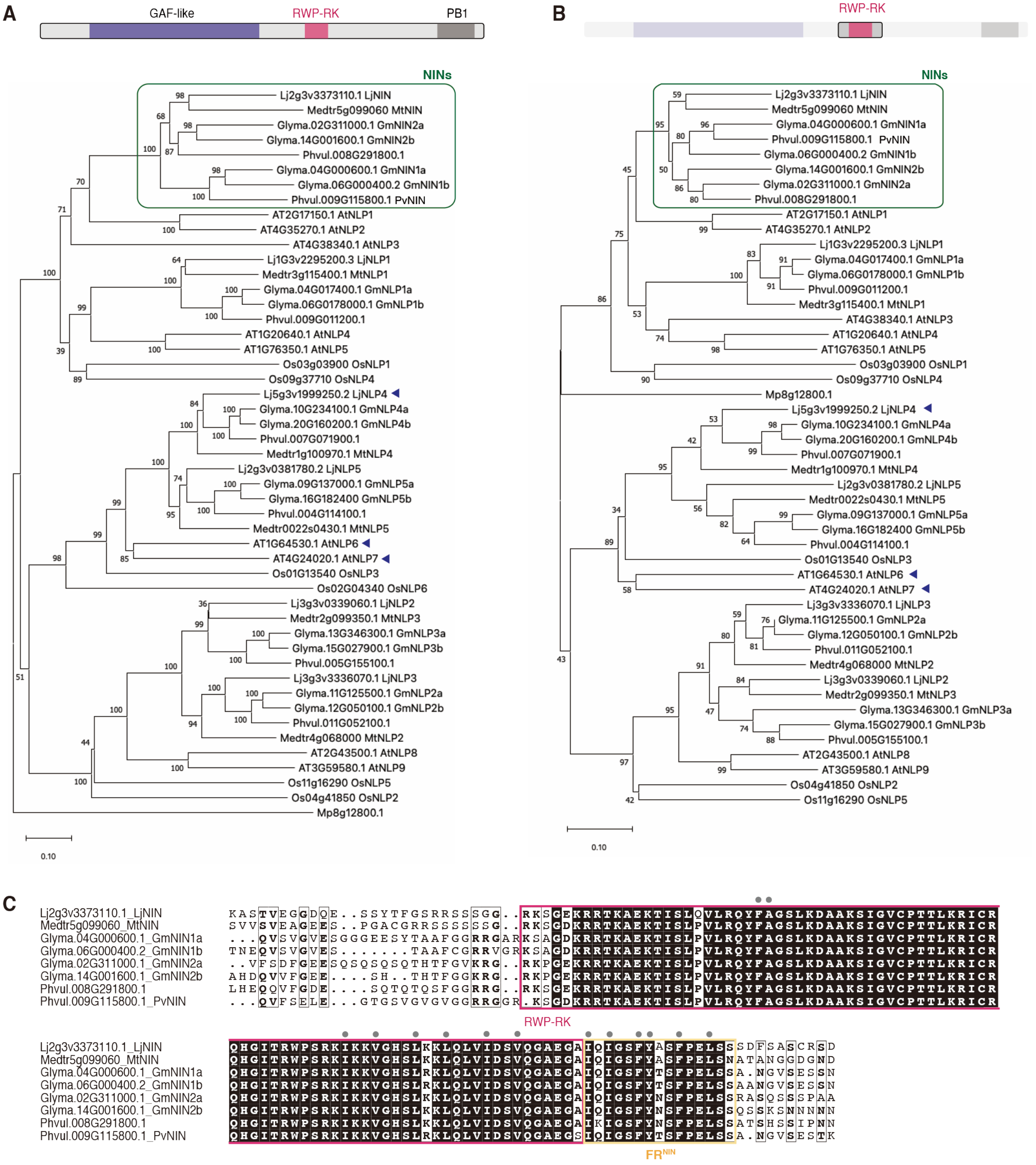


**Fig. S1.**

**Characterization of the DNA-binding regions from NINs.** (**A** and **B**) Phylogenetic trees of the NIN/NLP family proteins from *Lotus japonicus* (Lj), *Medicago truncatula* (Mt), *Glycine max* (Gm), *Phaseolus vulgari*s (Pv), *Arabidopsis thaliana* (At), *Oryza sativa* (Os), and *Marchantia polymorpha* (Mp). Full-length proteins (A) and DNA-binding regions (the RWP-RK DNA binding domains with 25 residues added before and after) (B) were compared and the trees were constructed by the neighbor-joining method. The percentages of replicate trees in which the associated taxa clustered together in the bootstrap test (1000 replicates) are shown next to the branches. Branch lengths are in the same units as those of the evolutionary distances. Accession numbers of the NIN/NLP family proteins used for the phylogenetic are as follows: Lj2g3v3373110.1 (LjNIN), Lj1g3v2295200.3 (LjNLP1), Lj3g3v0339060.1 (LjNLP2), Lj3g3v3336070.1 (LjNLP3), Lj5g3v1999250.2 (LjNLP4), and Lj2g3v0381780.2 (LjNLP5) from *L. japonicus*; Medtr5g099060 (MtNIN), Medtr3g115400.1 (MtNLP1), Medtr4g068000 (MtNLP2), Medtr2g099350.1 (MtNLP3), Medtr1g100970.1 (MtNLP4), Medtr0022s0430.1 (MtNLP5) from *M. truncatula*; Glyma.04G000600.1 (GmNIN1a), Glyma.06G000400.2 (GmNIN1b), Glyma.02G311000.1 (GmNIN2a), Glyma.14G001600.1 (GmNIN2b), Glyma.04G017400.1 (GmNLP1a), Glyma.06G0178000.1 (GmNLP1b), Glyma.11G125500.1 (GmNLP2a), Glyma.12G050100.1 (GmNLP2b), Glyma.13G346300.1 (GmNLP3a), Glyma.15G027900.1 (GmNLP3b), Glyma.10G234100.1 (GmNLP4a), Glyma.20G160200.1 (GmNLP4b), Glyma.09G137000.1 (GmNLP5a), Glyma.16G182400 (GmNLP5b) from *G. max*; Phvul.008G291800.1, Phvul.009G115800.1, Phvul.009G011200.1, Phvul.005G155100.1, Phvul.011G052100.1, Phvul.007G071900.1, Phvul.004G114100.1 from *P. vulgaris*; AT2G17150.1 (AtNLP1), AT4G35270.1 (AtNLP2), AT4G38340.1 (AtNLP3), AT1G20640.1 (AtNLP4), AT1G76350.1 (AtNLP5), AT1G64530.1 (AtNLP6), AT4G24020.1 (AtNLP7), AT2G43500.1 (AtNLP8), AT3G59580.1 (AtNLP9) from *A. thaliana*; Os03g03900 (OsNLP1), Os04g41850 (OsNLP2), Os01g13540 (OsNLP3), Os09g37710 (OsNLP4), Os11g16290 (OsNLP5), Os02g04340 (OsNLP6) from *O. sativa*; Mp8g12800.1 from *M. polymorpha*. OsNLP6 was excluded from the phylogenetic analyses of the DNA-binding regions because it has an incomplete RWP-RK domain. (**C**) A protein sequence alignment of the DNA-binding regions of the NINs from *L. japonicus*, *M. truncatula*, *G. max* and *P. vulgaris*. White boldface on black background and Black boldfaces on white background show the completely conserved and partially conserved amino-acid residues, respectively. The amino acid sequence identity within the RWP-RK domain among these NINs is 91%. Gray circles indicate the amino acid residues forming a dimer interface via hydrophobic and/or van der Waals interactions on the $FR^{LjNIN}$ in the Alphafold2-predicted structure.

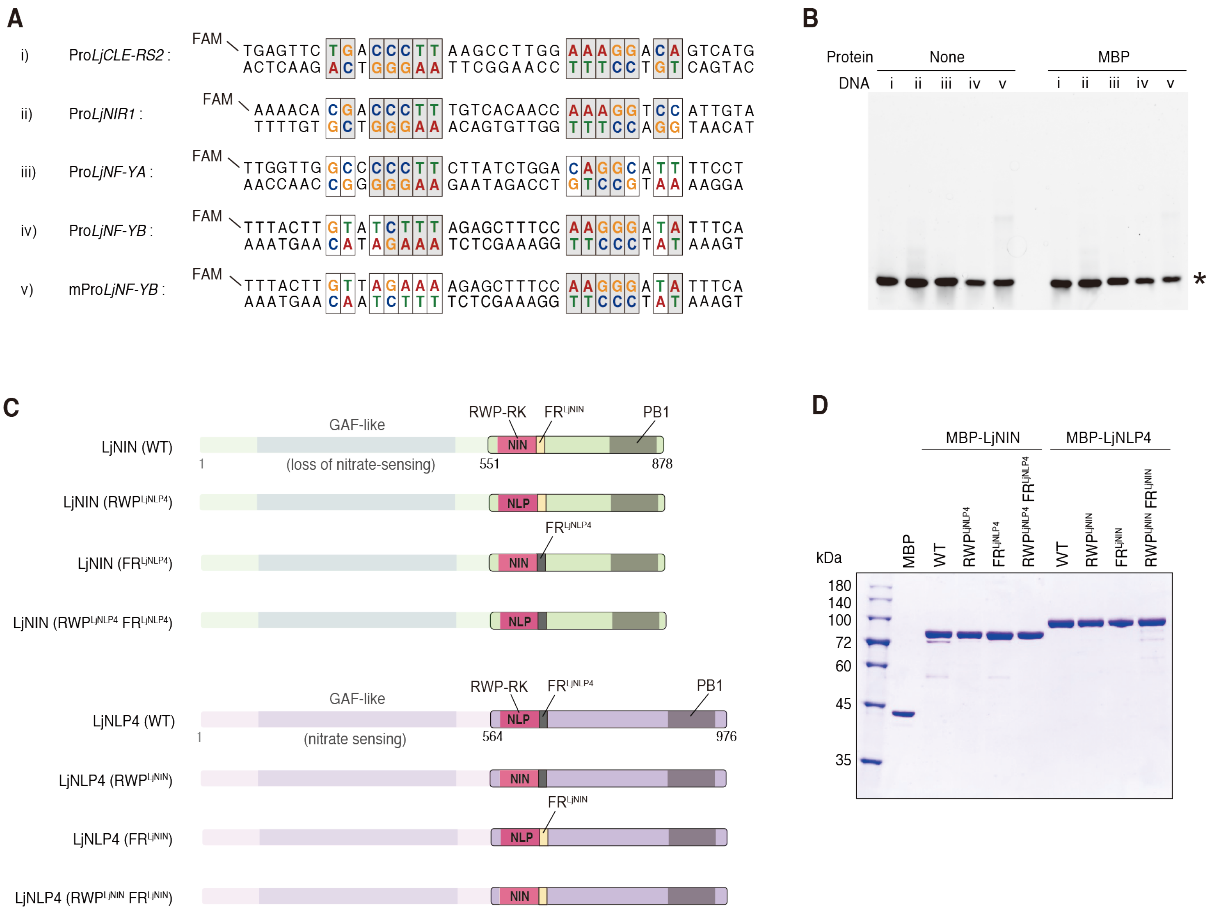


**Fig. S2.**

**DNA probes and the LjNIN/LjNLP proteins used for EMSA analyses.** (**A**) Complete sequences of the 5’ Carboxyfluorescein (FAM)-labeled DNA probes (i to v). (**B**) Control experiments of EMSA using 2.5 pmol DNA probes in the absence of protein (none) or in the presence of maltose-binding protein (MBP). The electrophoretic patterns are shown with an asterisk indicating the positions of free DNA. (**C**) The constructs of LjNIN (WT), LjNLP4 (WT), with the N-terminal region not involved in DNA binding deleted, and their chimeras with the RWP-RK and/or FR swapped. (**D**) SDS-PAGE analysis of the purified MBP and MBP-fused LjNIN/LjNLP proteins (related to Figs. 1D to G). The image shows CBB-stained proteins loaded onto gel in an amount of 10 pmol each.

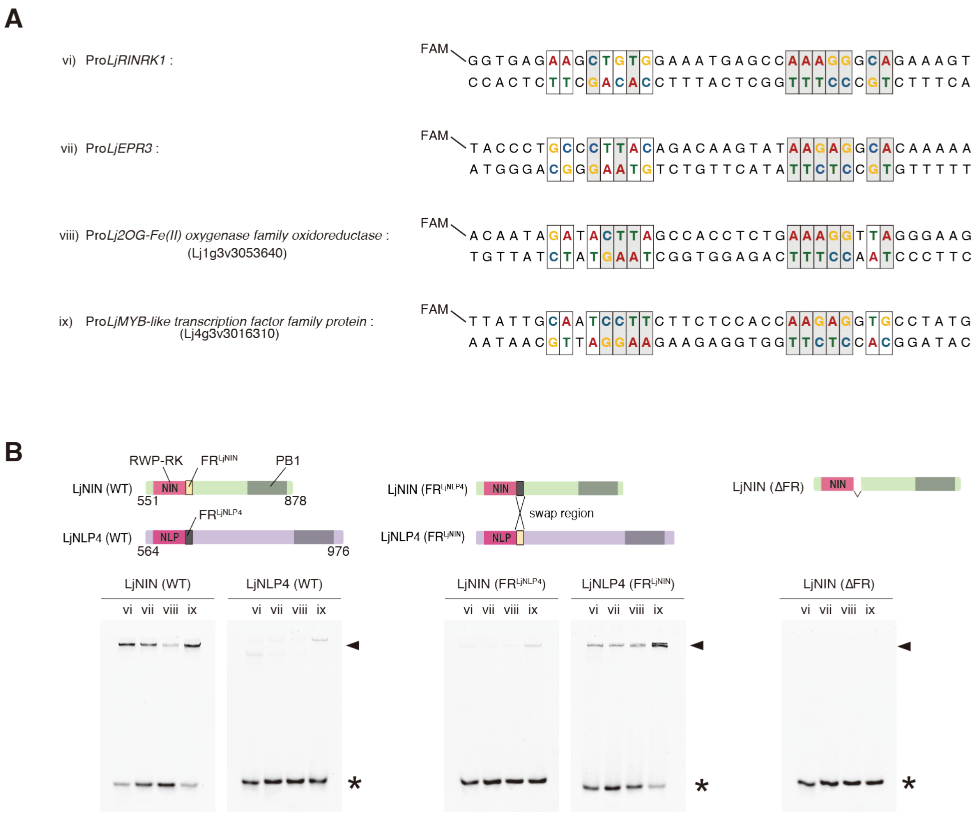


**Fig. S3.**

**Additional DNA-binding properties of LjNIN and LjNLP4 with or without FR$^{LjNIN}$.** (**A**) Additional FAM-labeled DNA probes (vi to ix) used for EMSA analyses. (**B**) EMSA results of LjNIN (WT), LjNLP4 (WT) proteins with the FR swapped (LjNIN (FR$^{LjNLP4}$) / LjNLP4 (FR$^{LjNIN}$)), and LjNIN protein with the FR deleted (LjNIN (ΔFR)) with the additional probes. The constructs of LjNIN/LjNLP4 proteins are shown at upper panels. Each recombinant protein fused to the MBP at N-terminus was reacted at 2 μM final conc. with 0.25 μM DNA probe (vi to ix). The electrophoretic patterns are shown at the lower panels with asterisks and arrowheads indicating the positions of free DNA and the protein-DNA complexes, respectively.

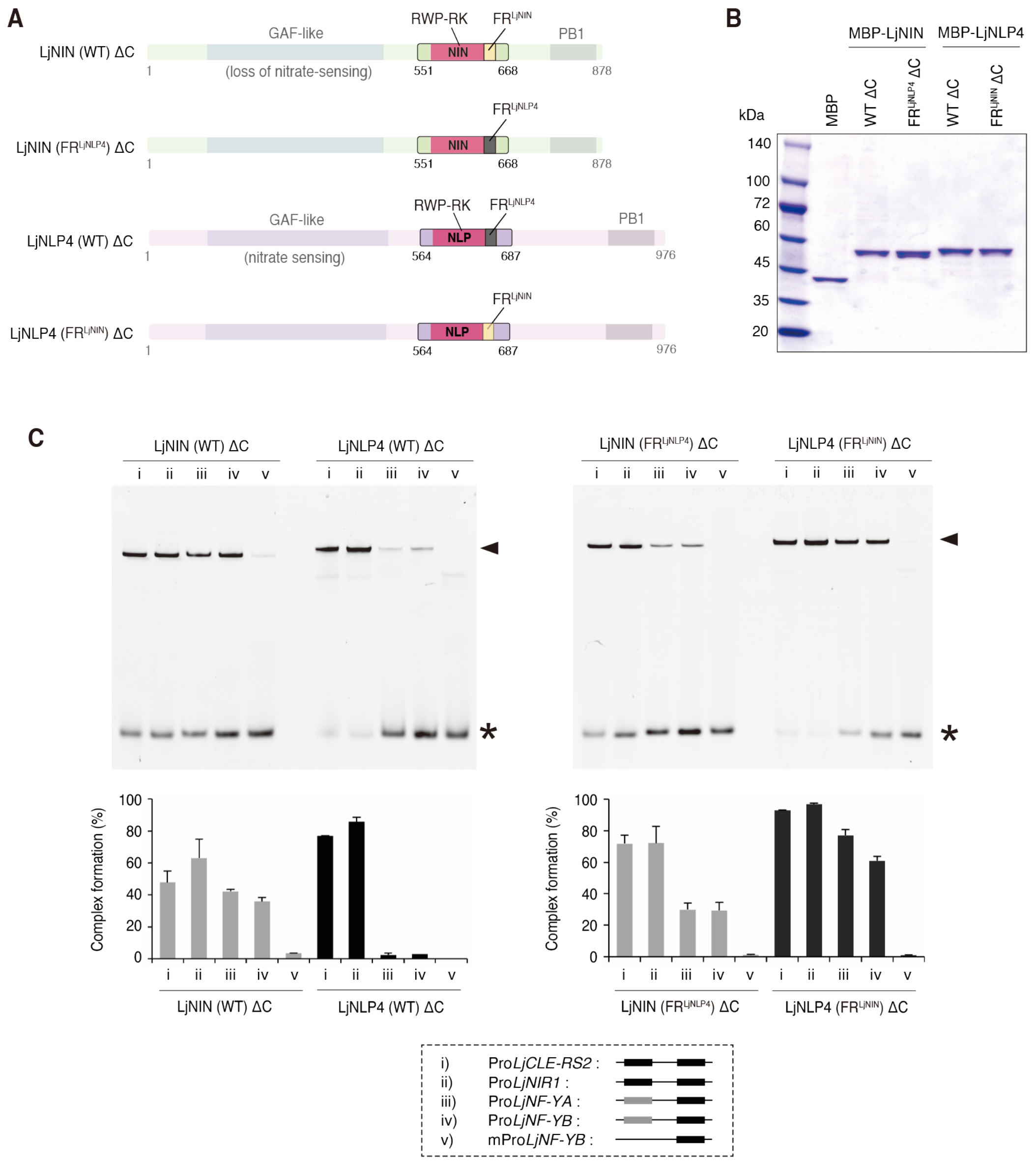


**Fig. S4.**

**DNA-binding properties of the DNA-binding modules of LjNIN and LjNLP4.** (A) The constructs of LjNIN (WT) ΔC, LjNLP4 (WT) ΔC, in which both the N-terminal and C-terminal regions were deleted, retaining the DNA-binding modules harboring the RWP-RK and FR. Chimeras with the FR swapped (LjNIN (FR$^{LjNLP4}$) ΔC/LjNLP4 (FR$^{LjNIN}$) ΔC) were also constructed. (B) SDS-PAGE analysis of the purified MBP and MBP-fused LjNIN/LjNLP4 ΔC proteins (related to Fig. 2, A to F). The image shows CBB-stained proteins loaded onto gel in an amount of 10 pmol each. (C) EMSA results of MBP-fused LjNIN/LjNLP ΔC proteins. Each protein was reacted at 2 µM final conc. with 0.25 µM DNA probe (i to v, shown in the lower panel). The electrophoretic patterns are shown at the upper panels with asterisks and arrowheads indicating the positions of free DNA and the protein-DNA complexes, respectively. Bar graphs at the middle panels showing the fluorescence densitometric profile are presented as mean ± s.e.m. (n=3).

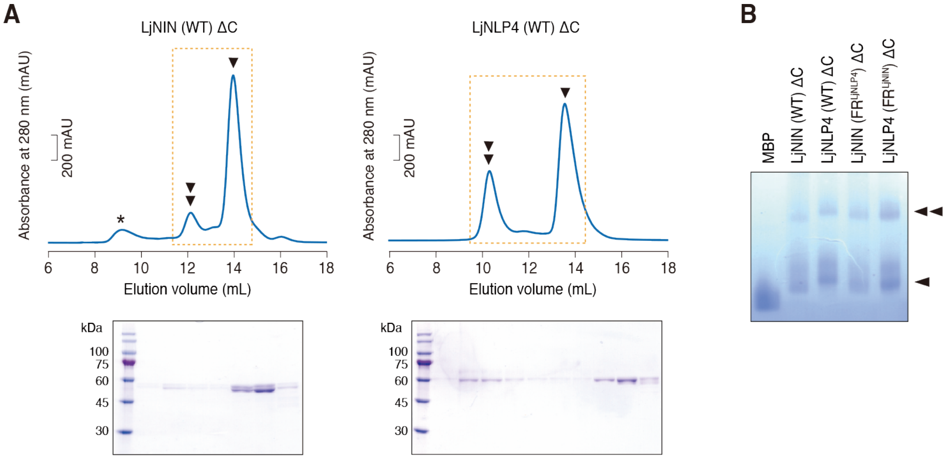


**Fig. S5.**

**Oligomeric analyses of the DNA-binding modules of LjNIN and LjNLP4.** (**A**) SEC analyses on the MBP-fused LjNIN (WT) ΔC and LjNLP4 (WT) ΔC. Chromatograms (upper panels) show monomeric and dimeric forms indicated by single and double arrowheads, respectively. The corresponding Coomassie-stained SDS-PAGE images (lower panels) were derived from the eluate fractions marked by dotted boxes in the chromatograms. Asterisk represents protein aggregation. (**B**) Blue-Native-PAGE results of MBP, MBP-fused LjNIN (WT) ΔC and LjNLP4 (WT) ΔC and their FR-swapped chimeras (LjNIN (FR$^{LjNLP4}$) ΔC and LjNLP4 (FR$^{LjNIN}$) ΔC). Each protein was loaded onto the gel at 2 µM final conc. A single arrowhead indicating the monomeric form and the double arrowheads indicating the dimeric form.

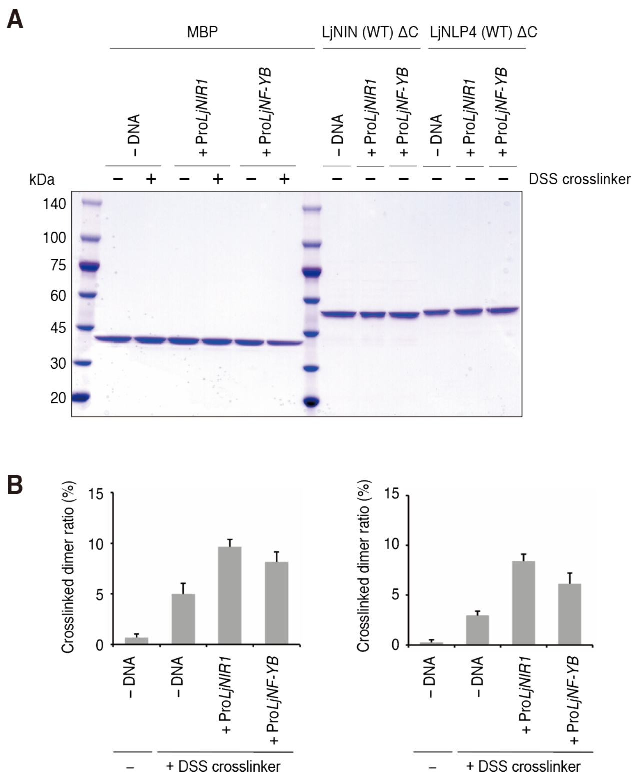


**Fig. S6.**

**Chemical crosslinking analyses of the DNA-binding modules of LjNIN and LjNLP4.** (**A**) Negative control experiments for crosslinking analysis using the amine-specific crosslinker DSS. SDS-PAGE results are shown for MBP alone, with or without DNA, under DSS-treated or DSS-free conditions, and for MBP-fused LjNIN (WT) ΔC and LjNLP4 (WT) ΔC in the presence of DNA under DSS-free conditions (corresponding to Fig. 2F). (**B**) Quantification of DSS-crosslinked dimer formation efficiency for LjNIN (WT) ΔC and LjNLP4 (WT) ΔC, with or without DNA. The proportion of DSS-crosslinked dimer species was quantified by densitometry of Coomassie-stained SDS-PAGE bands. Bar graphs show densitometric analysis of Coomassie-stained SDS-PAGE bands, presented as mean ± s.e.m. (n = 3). Corresponding raw SDS-PAGE gel images are shown in Fig. 2F. It should be noted that crosslinking efficiency depends not only on the efficiency of dimer formation, but also on the spatial proximity, accessibility, and flexibility of lysine residues and N-terminal amines.

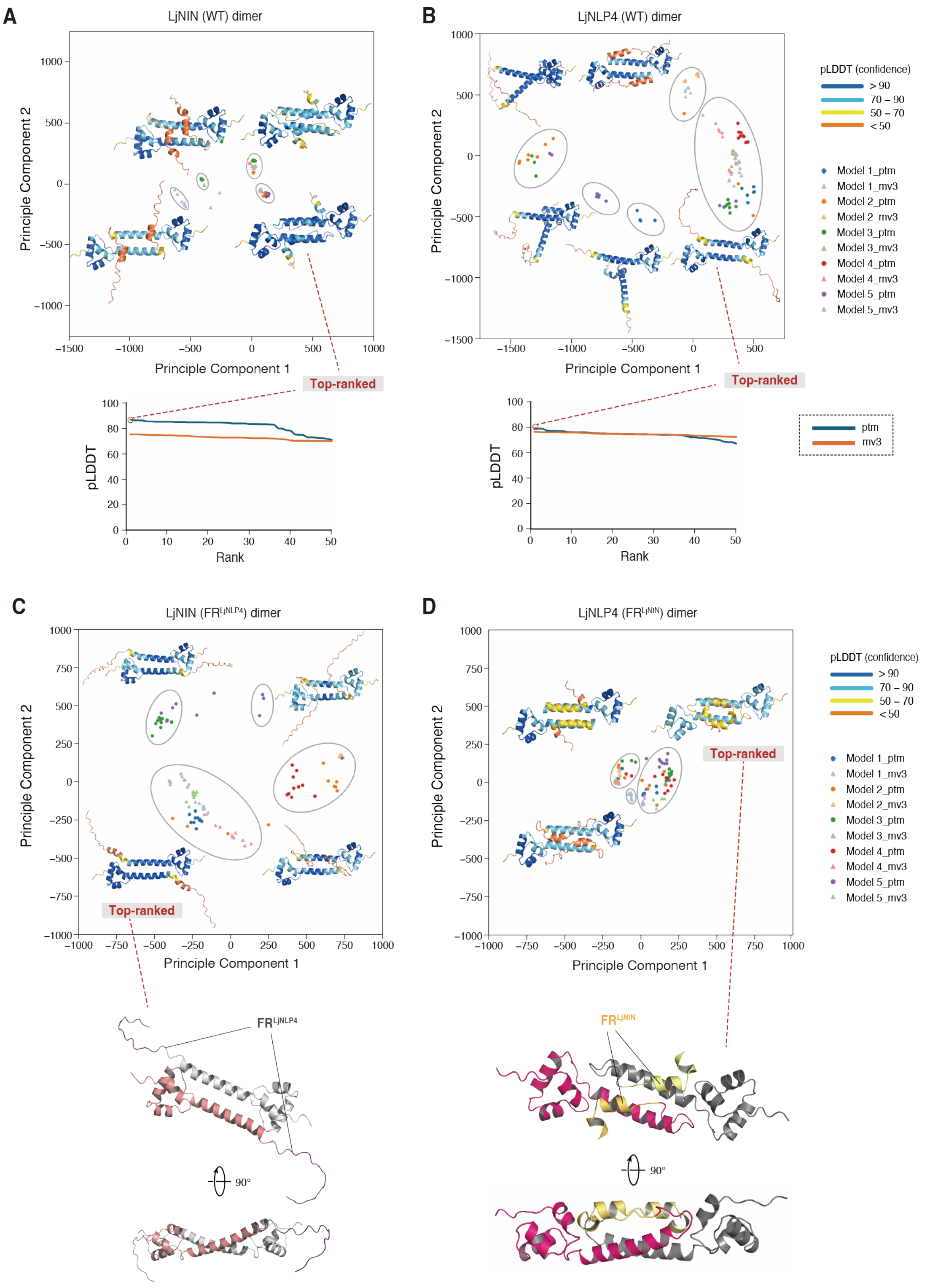


**Fig. S7.**

**Prediction of the LjNIN and LjNLP4 dimer structures by AlphaFold2.** (**A** and **B**) Clustering of structures of DNA-binding module dimers of the LjNIN (WT) (A) and LjNLP4 (WT) (B) predicted by alphafold2_monomer_ptm (ptm) and alphafold2_multimer_v3 (mv3) (upper panels), with pLDDT trend of the AlphaFold2 predicted structures of the LjNIN (WT) and the LjNLP4 (WT) dimers (bottom panels). Each fifty predicted structures by the two modes were outputted and mainly converged into several clusters. The predicted structures of LjNIN and LjNLP4 with the highest average value of pLDDT, which is the confidence of structure prediction, both belonged to the outputs from the alphafold2_monomer_ptm mode. In both top-ranked predicted structures, the average pLDDT score is approximately 80%, with most local pLDDT scores within the RWP-RK domains exceeding 90%. Most local pLDDT scores within $FR^{LjNIN}$ are above 70%. (**C** and **D**) Clustering of structures of DNA-binding module dimers of the LjNIN ($FR^{LjNLP4}$) (C) and the LjNLP4 ($FR^{LjNIN}$) (D) predicted by alphafold2_monomer_ptm (ptm) and alphafold2_multimer_v3 (mv3) (upper panels), with the top-ranked AlphaFold2 predicted structures of the LjNIN ($FR^{LjNLP4}$) the LjNLP4 ($FR^{LjNIN}$) dimers. Two different chains are depicted with different colors. The $FR^{LjNIN}$ are highlighted with yellow-based colors. In the top-ranked predicted structures of LjNIN ($FR^{LjNLP4}$), most local pLDDT scores within the RWP-RK domains exceed 90%, whereas those of LjNLP4 ($FR^{LjNIN}$) are relatively high as well, mostly above 70%.

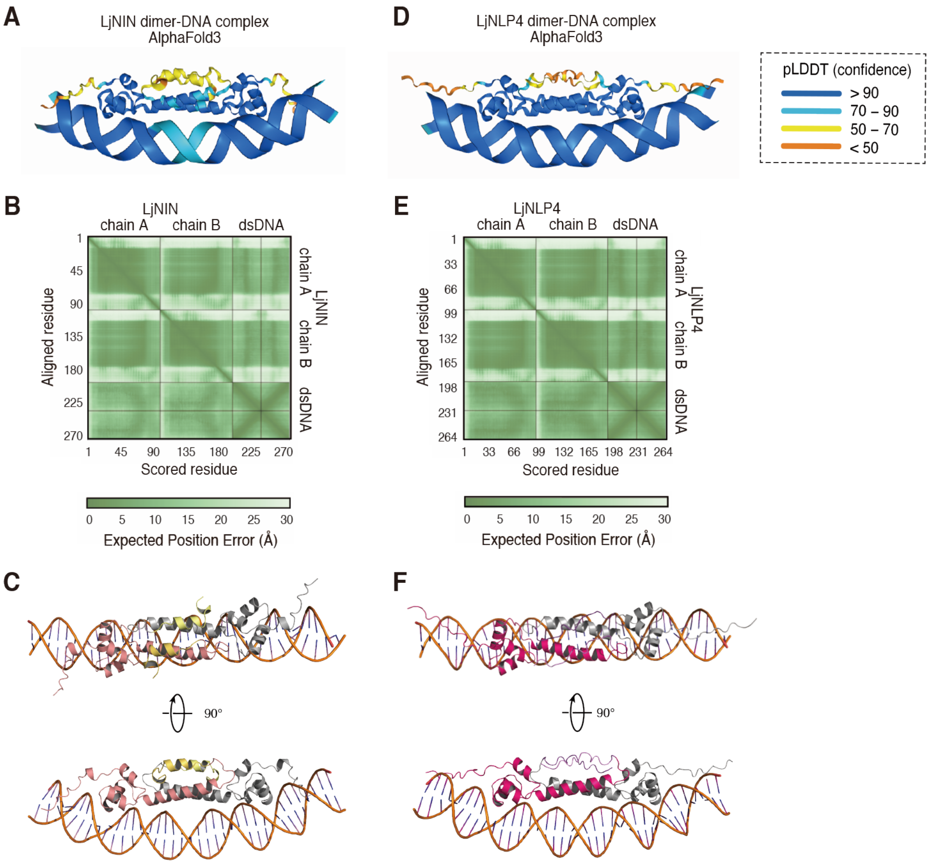


**Fig. S8.**

**AlpfaFold3 predicted structures of the LjNIN and LjNLP4 dimers in complex with DNA.** The top-ranked AlphaFold3-predicted structures of the DNA-binding module dimers of LjNIN (**A** to **C**) and LjNLP4 (**D** to **F**) in complex with target DNA (Pro*LjCLE-RS2* fragment). Structures are color coded by pLDDT scores (A and D), and the corresponding Predicted Aligned Error (PAE) values are shown (B and E). In both top-ranked predicted structures, most local pLDDT scores in the RWP-RK domains, and the LjNIN/LjNLP4-bound DNA exceed 90%, indicating high structural confidence in these regions. The relatively low PAE values (<10Å) between both the dimer subunits and the protein–double-stranded DNA (dsDNA) interfaces indicate high confidence in the predicted complex structures. Overall structures shown from two different views, with two different chains are depicted with different colors (C and F). The $\mathrm{FR}^{\mathrm{LjNIN}}$ are highlighted with yellow-based colors (C).

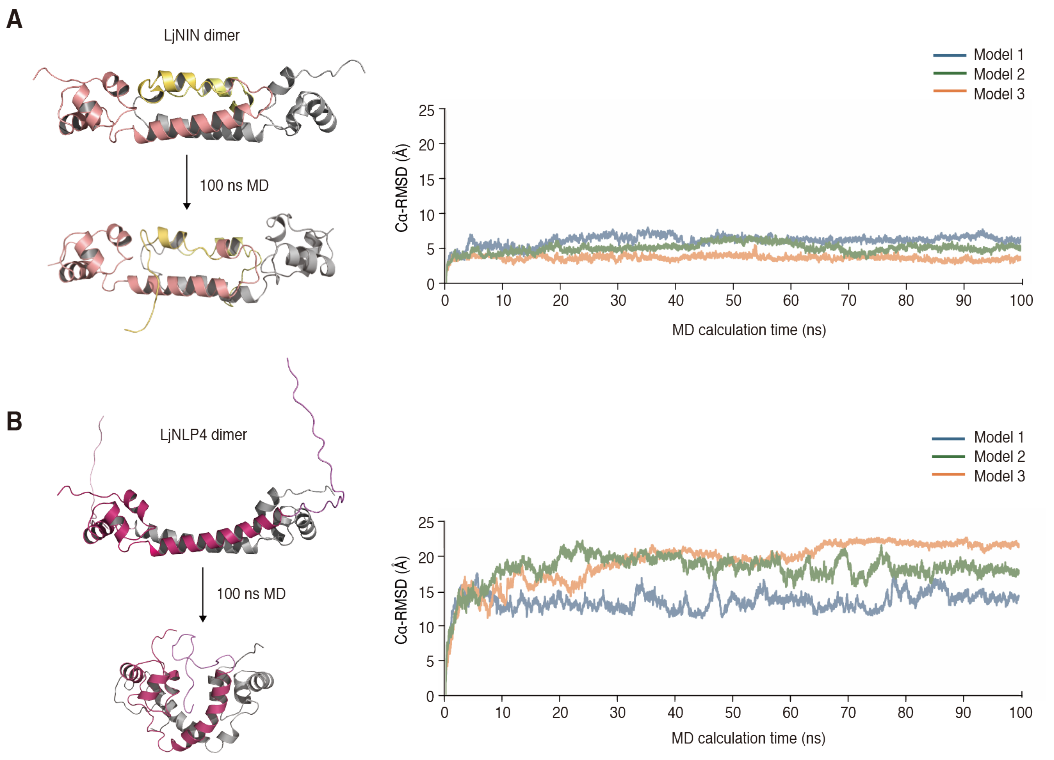


**Fig. S9.**

**MD simulations of predicted structures of the LjNIN and LjNLP4 dimers.** The top-ranked AlphaFold2 predicted structures of DNA-binding module dimers of LjNIN (**A**) and LjNLP4 (**B**) are used in three independent MD simulations for 100 ns. One representative structure after 100 ns simulation of three independent runs is shown for each in left panels. Each Cα RMSD chart within 100 ns is shown in right panels. LjNIN retained the dimeric structures optimal for DNA binding, as the root mean square deviation of main chains (Cα RMSD) converged to approximately 5Å, whereas LjNLP4 lost this conformation, with the Cα RMSD exceeding 15 Å.

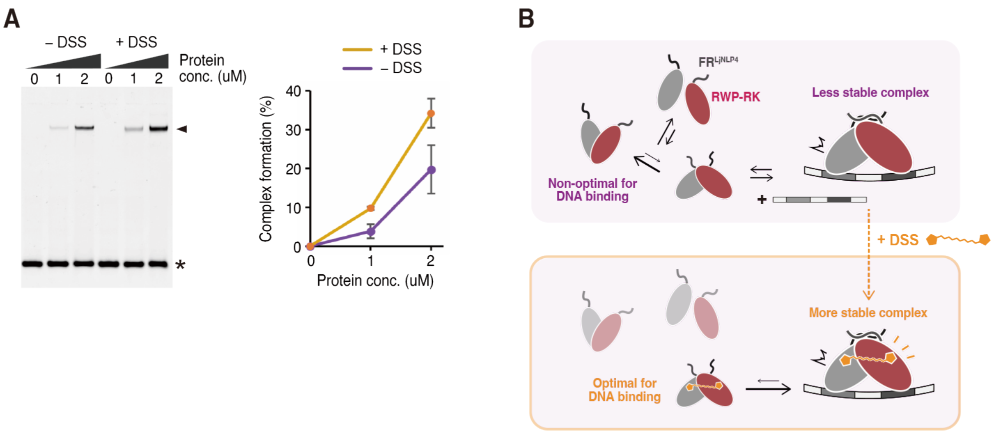


**Fig. S10.**

**Combination of chemical crosslinking and DNA-binding analyses for the DNA-binding module of LjNLP4.** (**A**) EMSA results of DSS-treated LjNLP4 (WT) ΔC. The protein, fused to MBP at the N-terminus, was reacted at final concentrations of 0, 1, and 2 µM with 0.25 µM of the Pro*LjNF-YB*-derived DNA probe, with or without DSS treatment. The electrophoretic patterns are shown at the left panel with asterisk and arrowhead indicating the positions of free DNA and the protein-DNA complexes, respectively. Bar graphs at the right panel showing the fluorescence densitometric profile are presented as mean ± s.e.m. (n=4). (**B**) Model of how DSS enhances the binding of the LjNLP4 DNA-binding module to Pro*LjNF-YB* containing unpreferred cis-elements. A fraction of LjNLP4 proteins is transiently recruited as dimers onto the DNA. Upon treatment with DSS, these dimers become covalently crosslinked into a conformation optimal for DNA binding. As a result, they are less likely to dissociate from the DNA, similar to the $FR^{NIN}$-stabilized dimers of LjNIN. Even after dissociating from DNA, the DSS-crosslinked LjNLP4 remains in a dimeric state that is optimal for binding, rather than reverting to a monomeric or non-optimal dimeric form, thereby enhancing overall DNA-binding efficiency.

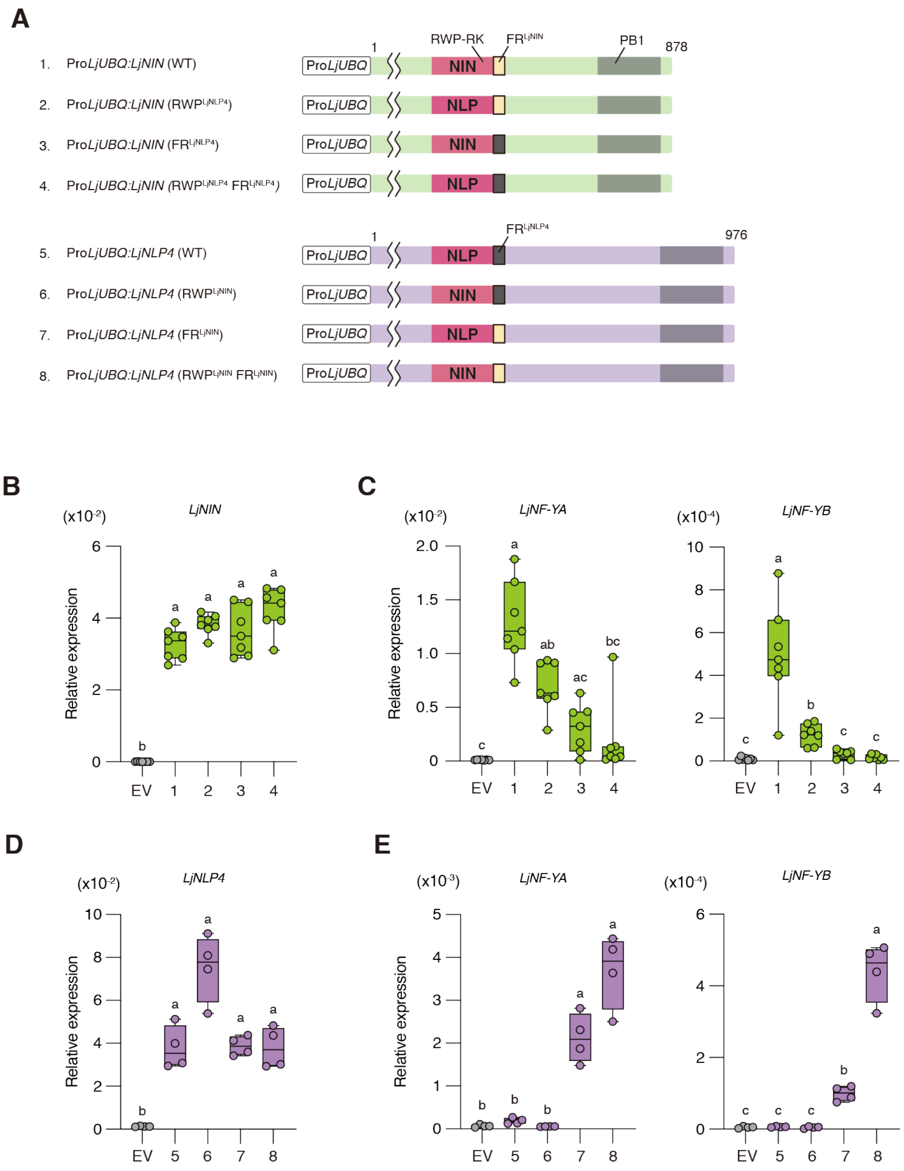


**Fig. S11.**

**Overexpression of the RWP-RK and the FR swapped LjNIN and LjNLP4 in *L. japonicus* hairy roots.** (**A**) The constructs of the RWP-RK and/or the FR swapped LjNIN/LjNLP4 proteins. (**B** to **E**) RT-qPCR analysis of *LjNIN*, *LjNLP4*, *LjNF-YA*, and *LjNF-YB* expressions in the transgenic hairy roots overexpressing the RWP-RK and/or the FR swapped *LjNIN* or *LjNLP4* by *LjUBQ* promoter. Transgenic plants were grown with 0 (B and C) or 0.5 mM $KNO_3$ (D and E) in the absence of rhizobia for 1 week. Numbers 1-8 correspond to the overexpressing constructs shown in (A). EV: empty vector (n = 7 or 4, each n contains hairy roots from 3 transgenic plants). Data were normalized by *LjUBQ* expression. Individual biological replicates are shown as dots. Different letters indicate statistically significant differences ($P < 0.05$, One-way ANOVA followed by multiple comparisons).

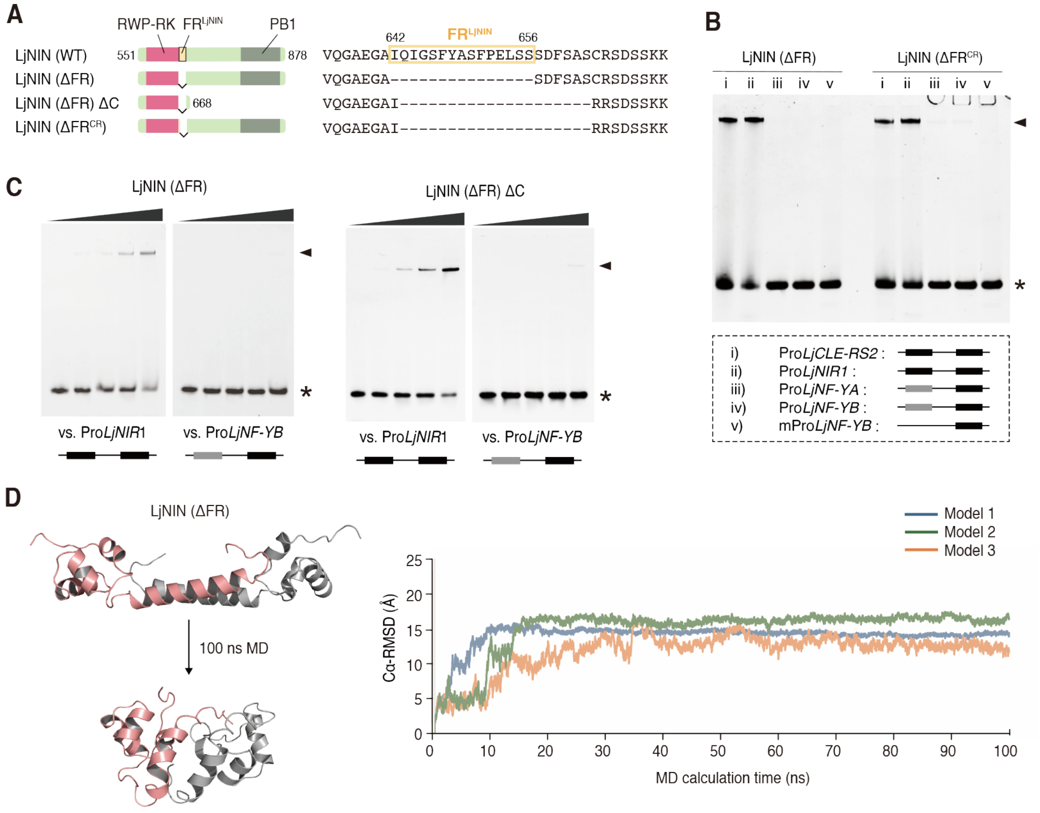


**Fig. S12.**

**Properties of LjNIN proteins with the FR deleted.** (**A**) The constructs of the LjNIN protein with the entire FR deleted (LjNIN (ΔFR)), its derivative lacking the C-terminal region (LjNIN (ΔFR) ΔC), and the LjNIN protein with the FR region deleted derived from ΔFR *nin* generated by CRISPR gene editing (LjNIN ($\Delta FR^{CR}$)). (**B**) EMSA results of LjNIN ($\Delta FR^{CR}$) compared with LjNIN (ΔFR). ΔFR protein fused to the MBP at N-terminus was reacted at 2 µM final conc. with 0.25 µM DNA probe (i to v; Fig. S2A). (**C**) EMSA results of LjNIN (ΔFR) and LjNIN (ΔFR) ΔC. Each ΔFR protein fused to the MBP at N-terminus were reacted at 0.25, 0.5, 1 and 2 µM final conc. The electrophoretic patterns are shown with an asterisk and an arrowhead. These experiments were repeated independently with similar results at least three times. Each protein producing a single shifted band regardless of protein concentration. (**D**) MD simulations of predicted structures of the LjNIN with the FR deleted in the DNA-free state. The top-ranked AlphaFold2 predicted structure of the LjNIN dimer with the FR deleted was used in three independent MD simulations for 100 ns. One representative structure after 100 ns simulation of three independent runs is shown for each in the left panel. The Cα RMSD chart within 100 ns is shown in the right panel. The Cα RMSD of LjNIN (ΔFR) converged above 15 Å, similar to LjNLP4 (WT) (Fig 2M, fig. S9B), but in contrast to LjNIN (WT), which remained stable at approximately 5 Å (Fig 2L, fig. S9A).

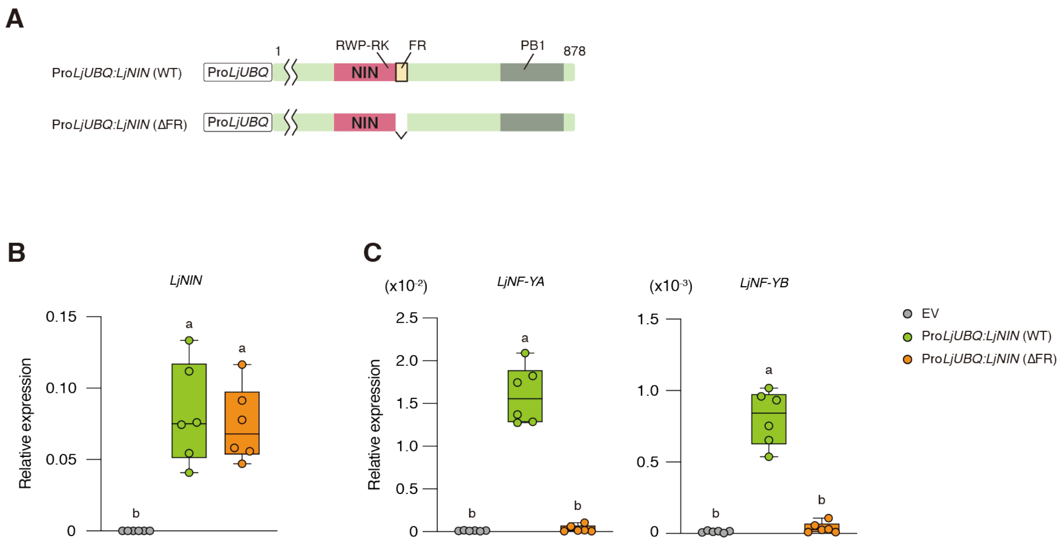


**Fig. S13.**

**Overexpression of LjNIN protein with the FR deleted in *L. japonicus* hairy roots.** (**A**) The constructs of LjNIN protein with the FR deleted (LjNIN (ΔFR)). (**B** and **C**) RT-qPCR analysis of *LjNIN*, *LjNF-YA*, and *LjNF-YB* expressions in the transgenic hairy roots overexpressing *LjNIN* or *LjNIN* (Δ*FR*) by *LjUBQ* promoter. Transgenic plants were grown in the absence of rhizobia for 1 week. EV: empty vector (n = 6, each n contains hairy roots from 3 transgenic plants). Data were normalized by *LjUBQ* expression. Individual biological replicates are shown as dots. Different letters indicate statistically significant differences (P < 0.05, One-way ANOVA followed by multiple comparisons).

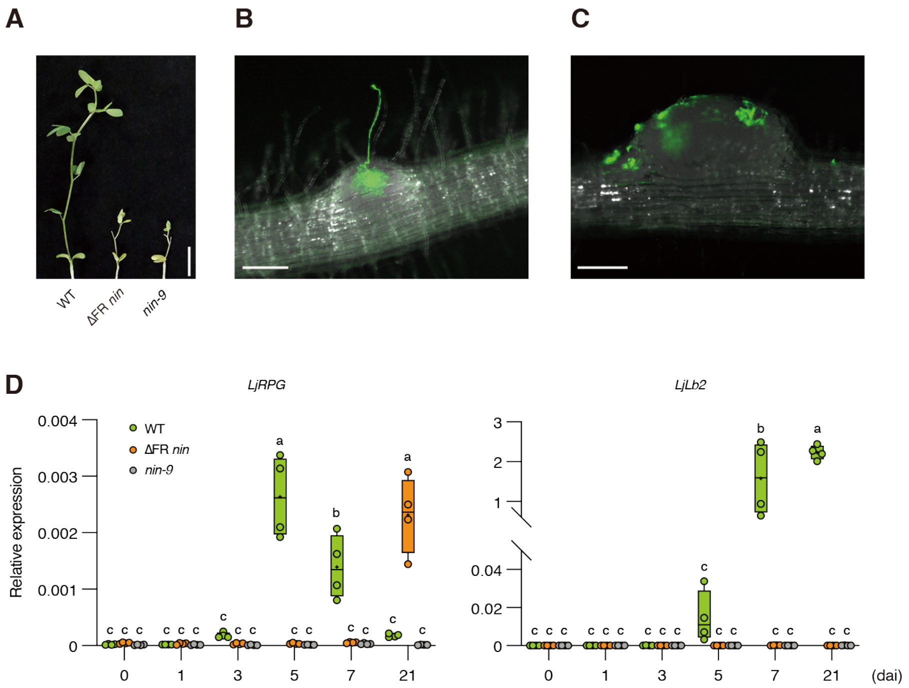


**Fig. S14.**
**Additional phenotypes of ΔFR *nin* mutants.** (**A**) Shoot phenotypes of WT, ΔFR *nin*, and *nin-9* plants at 28 dai. Bar, 1 cm. (**B** and **C**) Nodule primordia of WT at 7 dai (B) and ΔFR *nin* plants at 11 dai (C). Plants were inoculated with GFP-labelled rhizobia. (**D**) RT-qPCR analysis of *LjRPG* and *LjLb2* expressions in WT, ΔFR *nin*, and *nin-9* plants at 0, 1, 3, 5, 7, 21dai. Same cDNAs were used as those in Fig. 3R (n = 4, each n contains roots from at least 3 pants). Data were normalized by *LjUBQ* expression. Individual biological replicates are shown as dots. Different letters indicate statistically significant differences ($P < 0.05$, Two-way ANOVA followed by multiple comparisons). Bars, 1 cm (A), 100 µm [(B) and (C)].

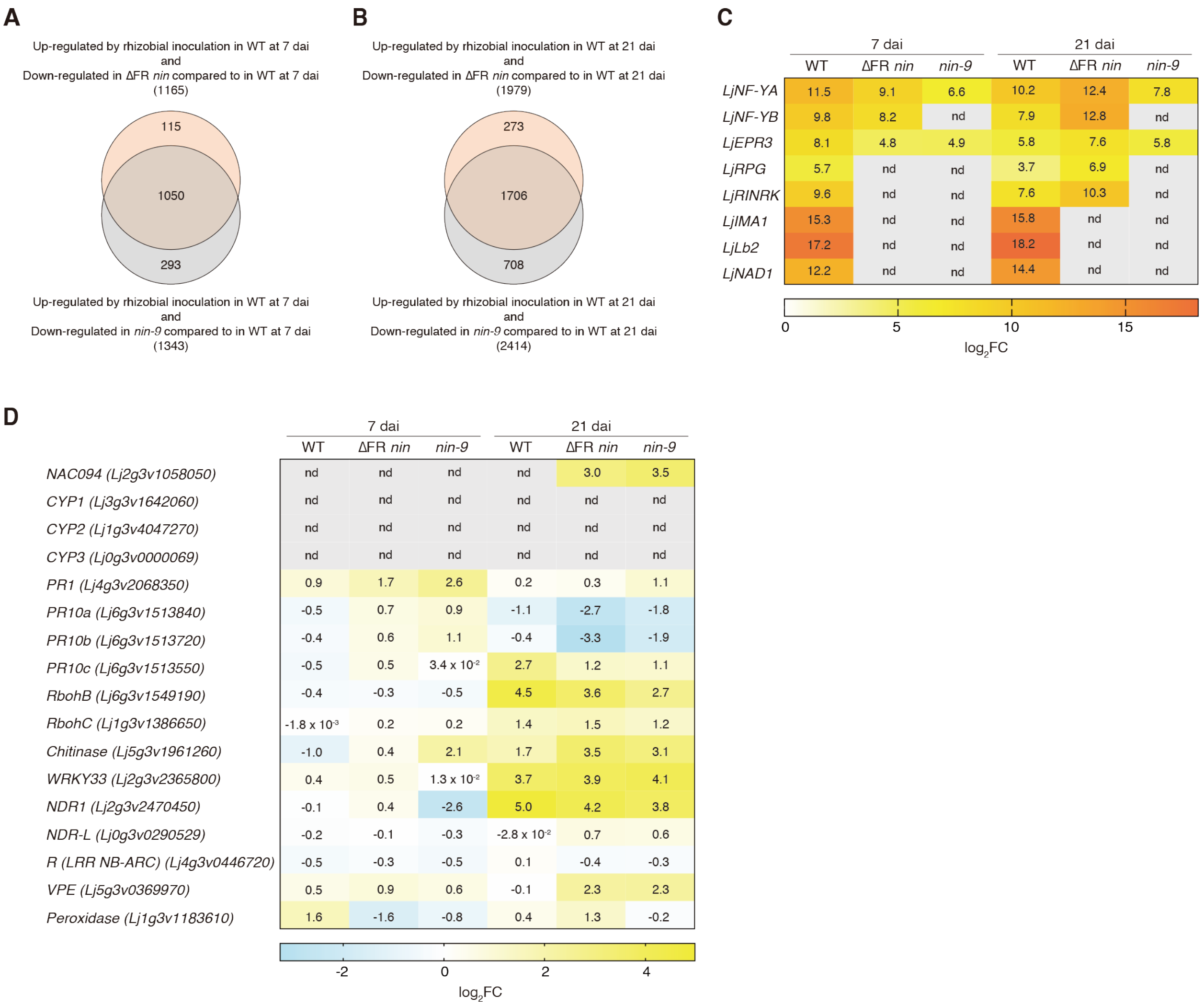


**Fig. S15.**

**RNA-seq analysis of ΔFR *nin* during root nodule symbiosis.** (**A** to **C**) Roots of WT, the ΔFR *nin* mutants, and the *nin-9* mutants at 0, 7, and 21 dai were used for transcriptome analysis. (n = 3, each n contains roots from at least 3 plants.) (A) Venn diagrams showing the overlap of the genes decreased expression between the ΔFR *nin* mutants and the *nin-9* mutants compared to WT. Genes upregulated by rhizobial inoculation in WT at 7 or 21 dai were identified based on a minimum 2-fold increase in expression ($\log_2$ fold change [$\log_2$FC] > 1, FDR < 0.05) relative to expression levels at 0 dai (non-inoculated) (Supplemental Data Set S1, A and D). Genes decreased expression in the ΔFR *nin* mutants or the *nin-9* mutants were identified based on similar criteria ($\log_2$FC < -1, FDR < 0.05), but relative to the expression levels in WT at 7 or 21 dai (Supplemental Data Set S1, B, C, E and F). (B and C) Relative expression levels of NIN-target genes (B) and defense or senescence marker genes (C). The numbers indicate the $\log_2$FC) relative to the expression level at 0 dai.

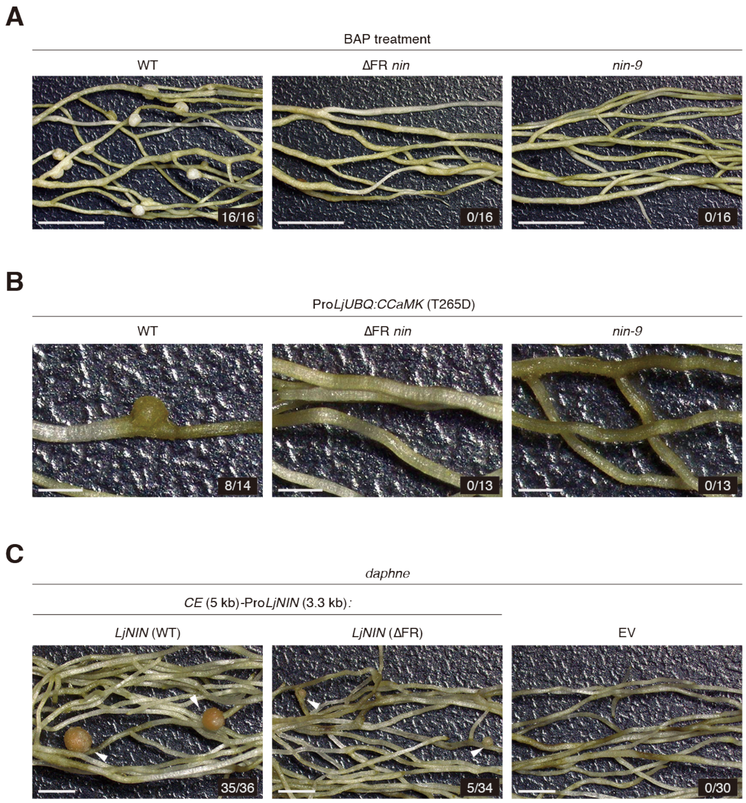


**Fig. S16.**

**The FR is necessary for the function of NIN in nodule organogenesis.** (**A** and **B**) Spontaneous nodule formations in ΔFR *nin* mutant induced by $10^{-7}$ M BAP treatment (A) or overexpressing constitutive active *CCaMK* (T265D) by *LjUBQ* promoter in hairy roots (B). Plants were grown with 0.5 (A) or 0.1 mM $KNO_3$ (B) in the absence of rhizobia for 6 weeks. (C) LjNIN (WT) or LjNIN protein with the FR deleted (LjNIN (ΔFR)) were expressed under *CE* (5 kb)-*LjNIN* promoter (3.3 kb) in hairy roots of *daphne* mutants. Transgenic plants were inoculated with rhizobia and nodules were observed at 28 dai. Numbers indicate the frequency of plants forming nodules (lower panels) among all tested plants. Bars, 5 mm (A), 1 mm (B), 2 mm (C).

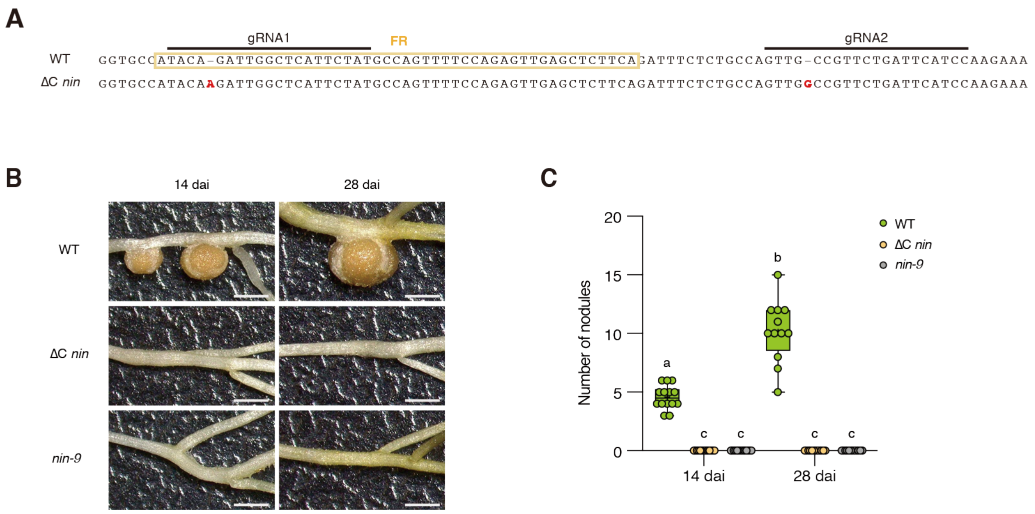


**Fig. S17.**
**Phenotypes of the *nin* mutant in which NIN is lacking the C-terminal region after RWP-RK.** (**A**) The location of the two base insertions causing a frameshift mutation in *nin* mutant in which NIN is lacking the C-terminal region after RWP-RK (ΔC *nin*) and gRNA regions used in CRISPR-Cas9 system. (**B** and **C**) Nodulation phenotypes of ΔC *nin* mutants at 14 and 28 dai. (B) Nodules. Bars, 1cm. (C) Number of total nodules. Different letters indicate statistically significant differences ($P < 0.05$, Two-way ANOVA followed by multiple comparisons).

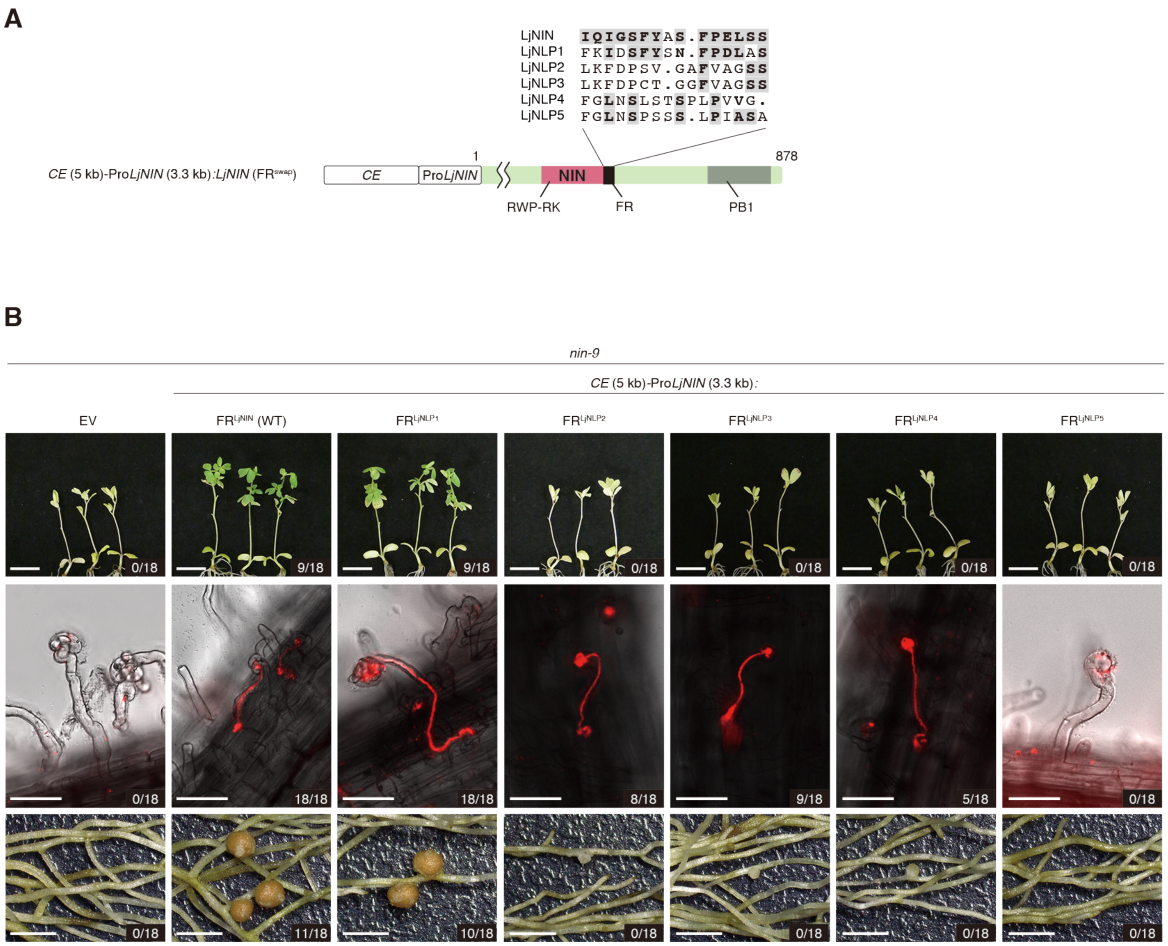


**Fig. S18.**

**The functional comparison of the $FR^{LjNLPs}$ and the $FR^{LjNIN}$.** (**A**) Protein sequence alignment of $FR^{LjNLPs}$. Highlighted letters on a gray background show the same amino acids as $FR^{LjNIN}$. Boldfaces on white background in protein sequence alignment of $FR^{LjNLPs}$ show amino acids with similar properties to those of $FR^{LjNIN}$. (**B**) Shoots (upper panels), ITs (middle panels) and nodules (lower panels) complemented by LjNIN proteins in which the FR was swapped with that of LjNLP1, LjNLP2, LjNLP3, LjNLP4, or LjNLP5 shown in (A). LjNIN ($FR^{swap}$) proteins were expressed under *CE* (5 kb)-*LjNIN* promoter (3.3 kb) in hairy roots of *nin-9* mutants. Transgenic plants were inoculated with DsRED-labelled rhizobia for 28 d. Numbers indicate the frequency of plants shoot recovered (upper panels), forming ITs (middle panels) or pink nodules (lower panels) among all transgenic plants. Bars, upper panels = 1 cm, middle panels = 50 µm, lower panels = 2 mm.

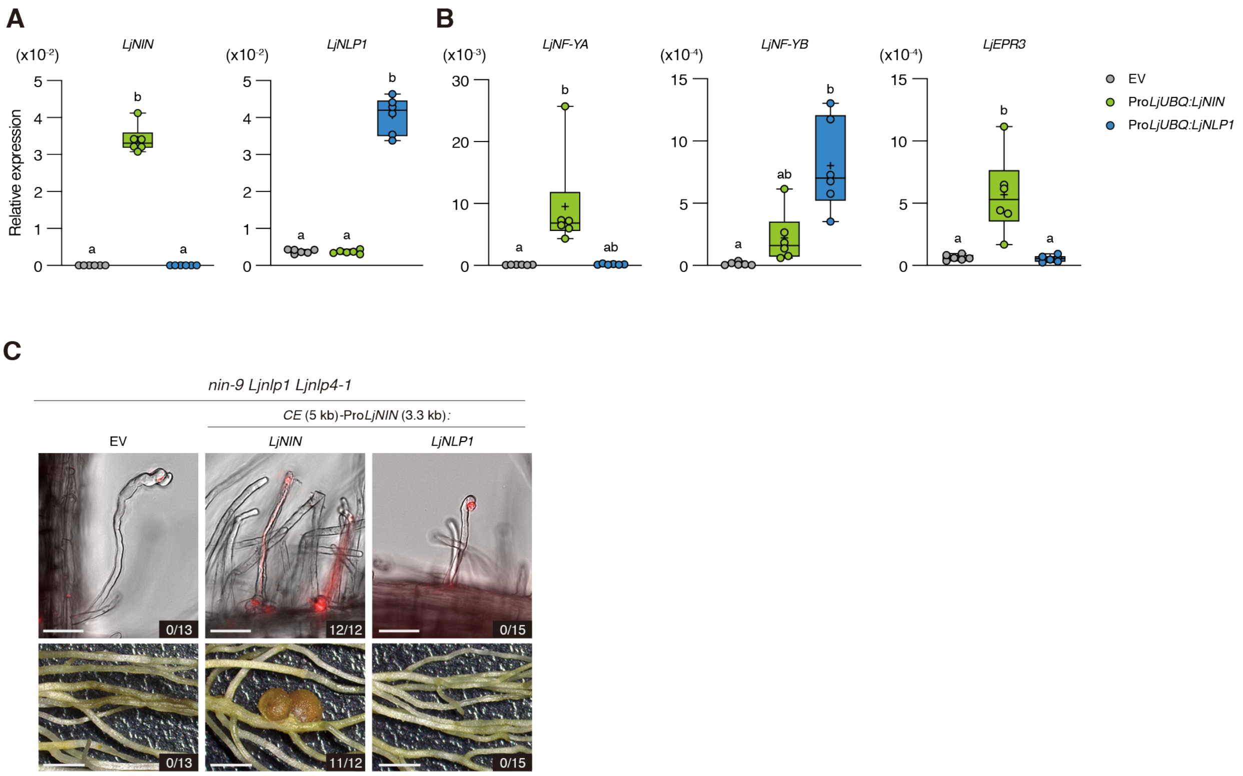


**Fig. S19.**

**Biological characterization of LjNLP1.** (**A** and **B**) RT-qPCR analysis of *LjNIN*, *LjNLP1*, *LjNF-YA*, *LjNF-YB*, and *LjEPR3* expressions in the transgenic hairy roots overexpressing *LjNIN* or *LjNLP1* by *LjUBQ* promoter. Transgenic plants were grown with 0.5 mM $KNO_3$ in the absence of rhizobia for 1 week. EV: empty vector (n = 6, each n contains hairy roots from 3 transgenic plants). Data were normalized by *LjUBQ* expression. Individual biological replicates are shown as dots. Different letters indicate statistically significant differences ($P < 0.05$, One-way ANOVA followed by multiple comparisons). (**C**) *LjNIN* or *LjNLP1* were expressed under *CE* (5 kb)-*LjNIN* promoter (3.3 kb) in hairy roots of *nin-9 Ljnlp1 Ljnlp4-1* triple mutants. Transgenic plants were inoculated with DsRED-labelled rhizobia in the presence of 5 mM $KNO_3$ for 28 d. ITs and nodules are shown at upper and lower panels respectively. Numbers indicate the frequency of plants forming ITs (upper panels) or nodules (lower panels) among all transgenic plants. Bars, upper panels = 50 µm, lower panels = 2 mm.

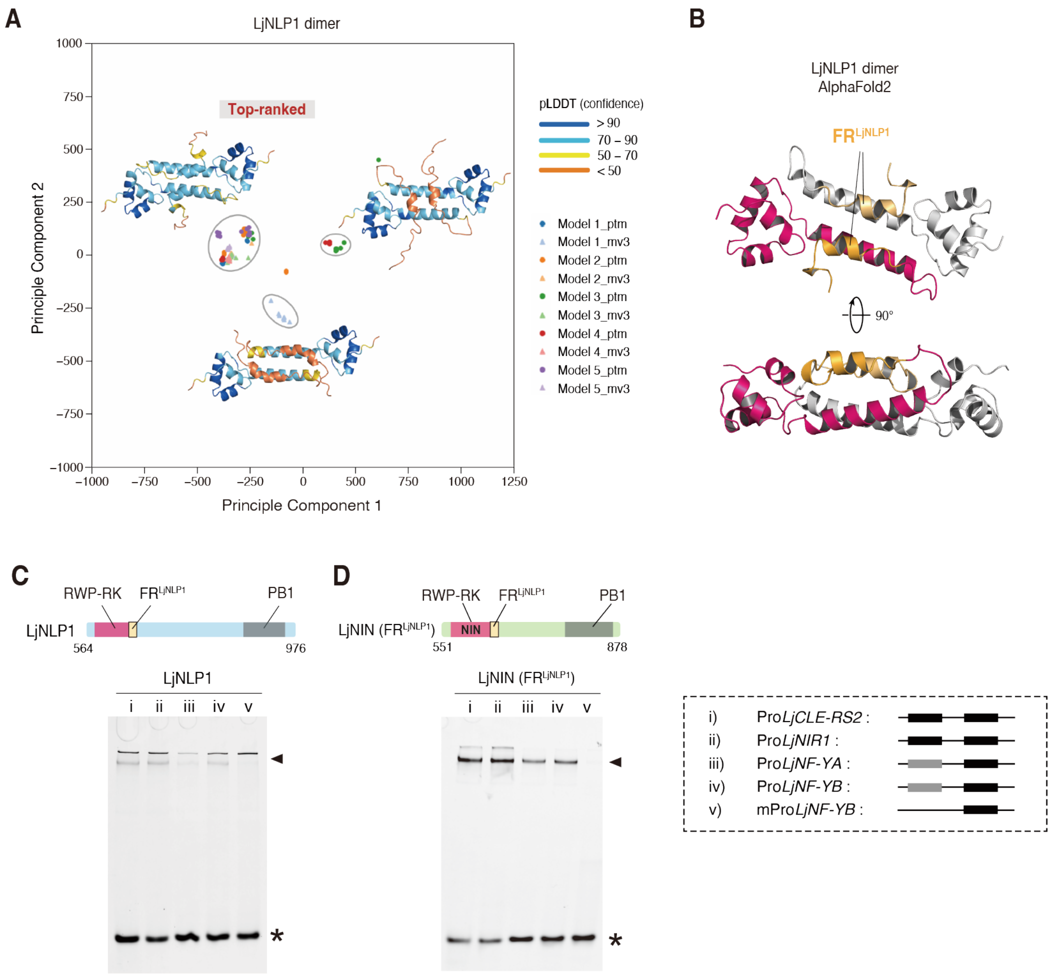


**Fig. S20.**

**Biochemical and structural characterization of LjNLP1.** (**A**) Clustering of structures of the DNA-binding module dimer of LjNLP1 predicted by alphafold2_monomer_ptm (ptm) and alphafold2_multimer_v3 (mv3). (**B**) Top-ranked AlphaFold2 predicted structure of the DNA-binding module dimer of the LjNLP1. Two different chains are depicted with different colors. The $FR^{LjNLP1}$ are highlighted with orange-based colors. In the top-ranked predicted structure, most local pLDDT scores within the RWP-RK and FR are above 70%. (**C** and **D**) EMSA results of LjNLP1 and LjNIN ($FR^{LjNLP1}$). The constructs of the NIN/NLP proteins are shown at the upper panels. Each recombinant protein fused to the MBP at N-terminus was reacted at 2 µM final conc. with 0.25 µM DNA probe (i to v). The electrophoretic patterns are shown at the lower panels with asterisks and arrowheads indicating the positions of free DNA and the protein-DNA complexes, respectively.

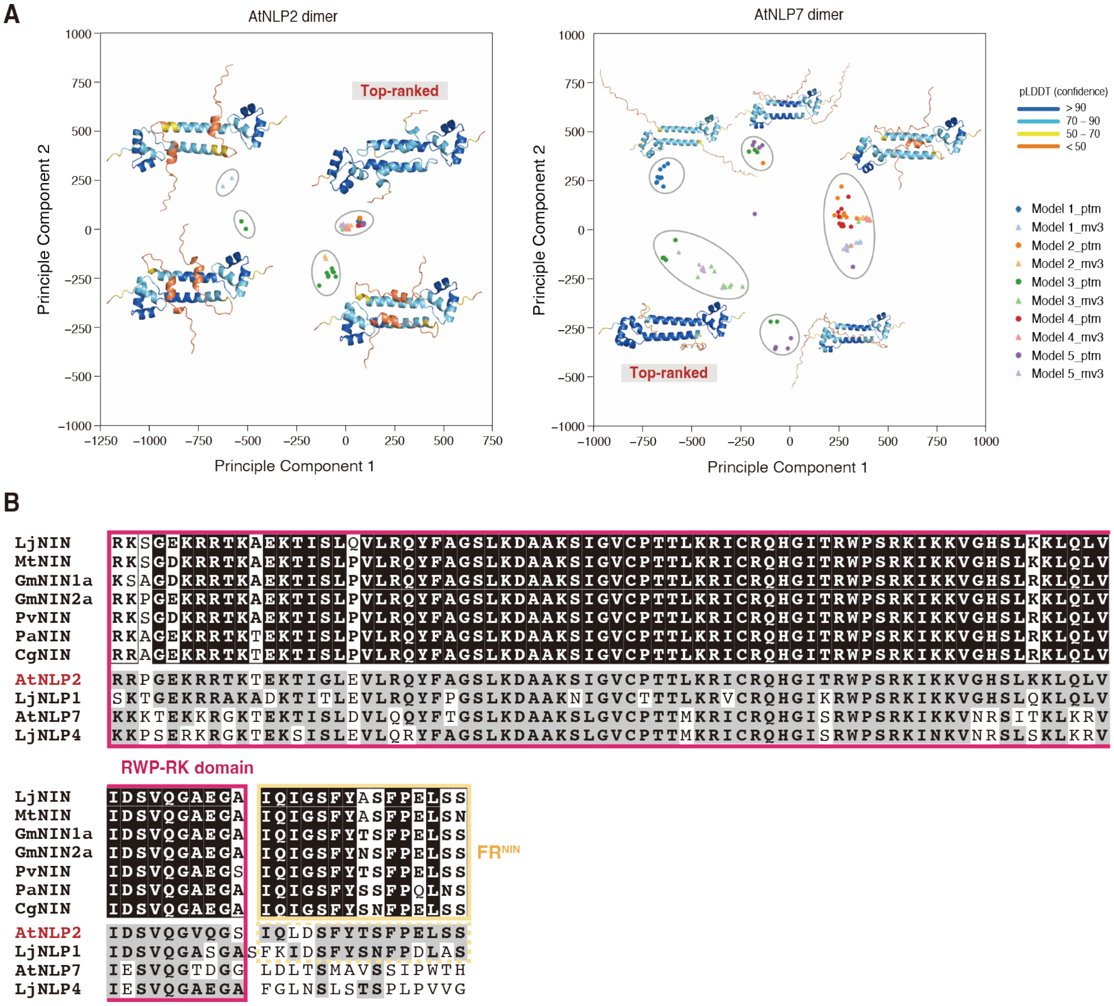


**Fig. S21.**

**Predicted structures and amino-acid sequences of AtNLP2 and other NIN/NLPs.** (**A**) Clustering of structures of the DNA-binding module dimers of AtNLP2 and AtNLP7 predicted by alphafold2_monomer_ptm (ptm) and alphafold2_multimer_v3 (mv3). In both top-ranked predicted structures, most local pLDDT scores within the RWP-RK domains are exceeding 90%. Most local pLDDT scores within $FR^{LjNIN}$ are above 70%. (**B**) Amino-acid sequences of the DNA-binding modules of the NLP/NINs harboring the RWP-RK and FR. The sequences include NINs from *L. japonicus* (Lj), *M. truncatula* (Mt), *G. max* (Gm), *P. vulgaris* (Pv), *P. andersonii* (Pa), and *C. glauca* (Cg); AtNLP2 and AtNLP7 from *A. thaliana* (At); and LjNLP1 and LjNLP4 from *L. japonicus* (Lj). In the NIN alignment, white bold letters on a black background indicate fully conserved amino-acid residues, while black bold letters on a white background indicate partially conserved residues. In the sequences of AtNLP2, AtNLP7, LjNLP1, and LjNLP4, highlighted letters on a gray background indicate residues identical to the corresponding residues in NINs.

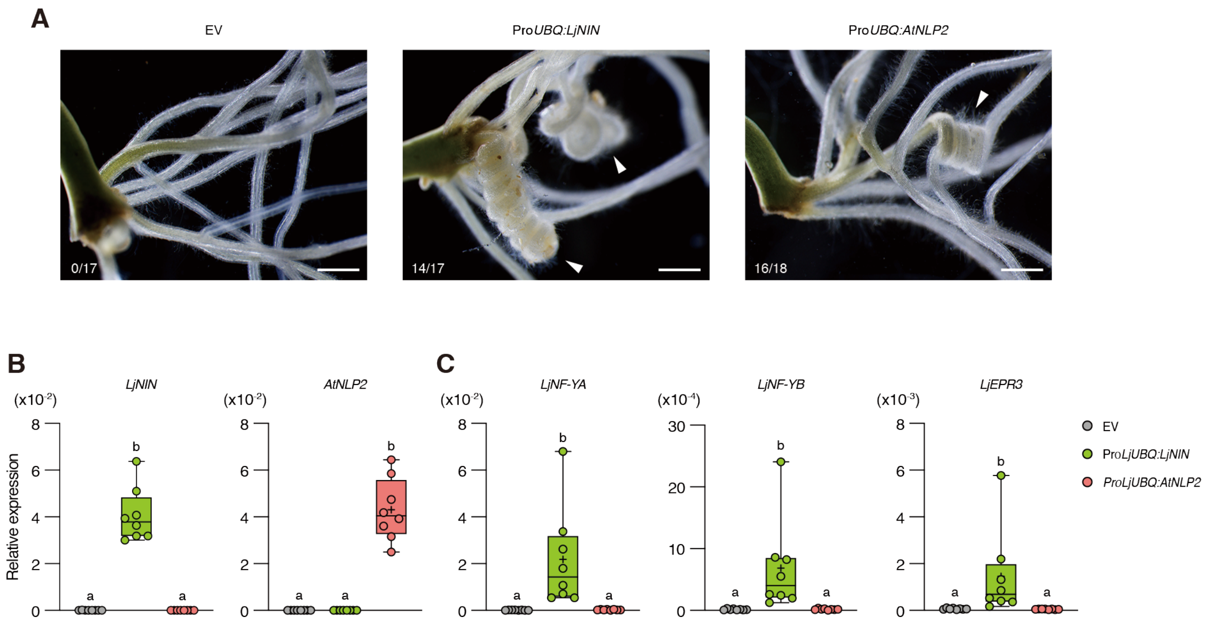


**Fig. S22.**
**Overexpression of *AtNLP2* in *L. japonicus* hairy root.** (**A**) Phenotypes of transgenic hairy roots overexpressing *LjNIN* or *AtNLP2* by *LjUBQ* promoter. Transgenic plants were grown with 0.5 mM $KNO_3$ in the absence of rhizobia for 1 week. Arrowheads indicate abnormal roots characterized by excessive helical structures which were often observed in *LjNIN* overexpressing hairy roots. Numbers indicate the frequency of plants forming abnormal roots among all transgenic plants. Bars =1 mm. (**B** and **C**) RT-qPCR analysis of *LjNIN*, *AtNLP2*, *LjNF-YA*, *LjNF-YB*, and *LjEPR3* expressions in the transgenic hairy roots overexpressing *LjNIN* or *AtNLP2* by *LjUBQ* promoter. Transgenic plants were grown without $KNO_3$ in the absence of rhizobia for 1 week. (n = 8, each n contains hairy roots from 3 transgenic plants). Data were normalized by *LjUBQ* expression. Individual biological replicates are shown as dots. Different letters indicate statistically significant differences ($P < 0.05$, One-way ANOVA followed by multiple comparisons).

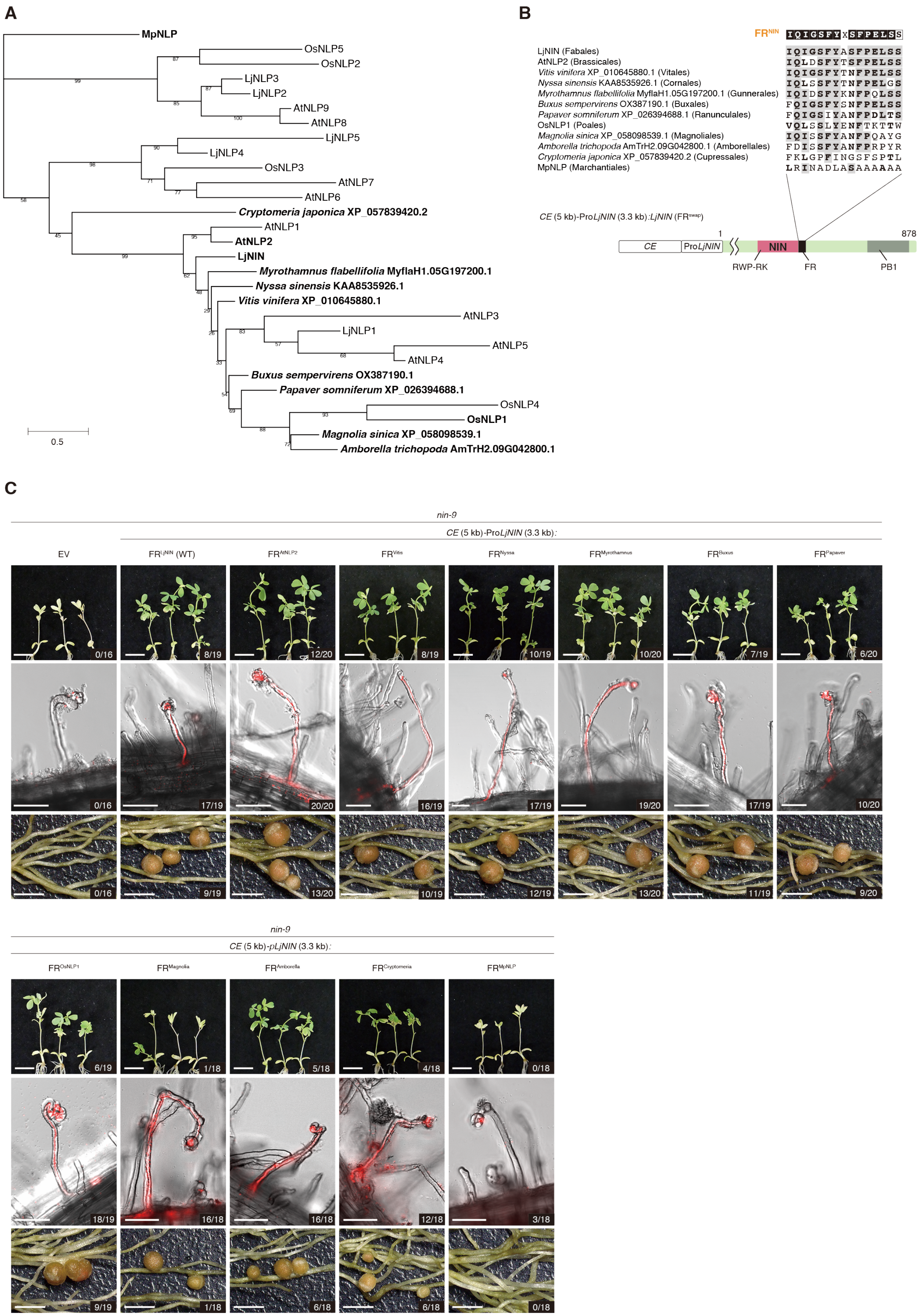
A
MpNLP
OsNLP5
OsNLP2
LjNLP3
LjNLP2
AtNLP9
AtNLP8
LjNLP5
LjNLP4
OsNLP3
AtNLP7
AtNLP6
Cryptomeria japonica XP_057839420.2
AtNLP1
AtNLP2
LjNIN
Myrothamnus flabellifolia MyflaH1.05G197200.1
Nyssa sinensis KAA8535926.1
Vitis vinifera XP_010645880.1
AtNLP3
LjNLP1
AtNLP5
AtNLP4
Buxus sempervirens OX387190.1
Papaver somniferum XP_026394688.1
OsNLP4
OsNLP1
Magnolia sinica XP_058098539.1
Amborella trichopoda AmTrH2.09G042800.1
0.5
B
FRNIN IQIGSFYxSFPELSS
LjNIN (Fabales) IQIGSFYASFPELSS
AtNLP2 (Brassicales) IQLDSFYTSFPELSS
Vitis vinifera XP_010645880.1 (Vitales) IQIGSFYTNFPELSS
Nyssa sinensis KAA8535926.1 (Cornales) IQLSSFYTNFPELGS
Myrothamnus flabellifolia MyflaH1.05G197200.1 (Gunnerales) IQIDSFYKNFPQLSS
Buxus sempervirens OX387190.1 (Buxales) FQIGSFYSNFPELSS
Papaver somniferum XP_026394688.1 (Ranunculales) FQIGSIYANFPDLTS
OsNLP1 (Poales) VQLSSLYENFTKTTW
Magnolia sinica XP_058098539.1 (Magnoliales) IQISSFYANFPQAYG
Amborella trichopoda AmTrH2.09G042800.1 (Amborellales) FDISSFYANFPRPYR
Cryptomeria japonica XP_057839420.2 (Cupressales) FKLGPFINGSFSPTL
MpNLP (Marchantiales) LRINADLASAAAAAA
CE (5 kb)-ProLjNIN (3.3 kb):LjNIN (FRswap)
1
878
CE
ProLjNIN
NIN
RWP-RK
FR
PB1
C
nin-9
CE (5 kb)-ProLjNIN (3.3 kb):
EV
FRLjNIN (WT)
FRAtNLP2
FRVitis
FRNyssa
FRMyrothamnus
FRBuxus
FRPapaver
0/16
8/19
12/20
8/19
10/19
10/20
7/19
6/20
0/16
17/19
20/20
16/19
17/19
19/20
17/19
10/20
0/16
9/19
13/20
10/19
12/19
13/20
11/19
9/20
nin-9
CE (5 kb)-pLjNIN (3.3 kb):
FROsNLP1
FRMagnolia
FRAmborella
FRCryptomeria
FRMpNLP
6/19
1/18
5/18
4/18
0/18
18/19
16/18
16/18
12/18
3/18
9/19
1/18
6/18
6/18
0/18

**Fig. S23.**

**Functional analysis of FR of NLPs from land plant species.** (**A**) Phylogenetic tree was constructed using DNA-binding regions (the RWP-RK DNA-binding domains with 25 residues added before and after) of LjNIN and its closest homologous NLPs from eudicots (*A. thaliana* (At), *Vitis vinifera*, *Nyssa sinensis*, *Myrothamnus flabellifolius*, *Buxus sempervirens*, *Papaver somniferum*), monocot (*O. sativa* (Os)), magnoliid (*Magnolia sinica*), basal angiosperm (*Amborella trichopoda*), gymnosperm (*Cryptomeria japonica*), and moss (*M. polymorpha* (Mp)) as well as all NLPs from *L. japonicus*, *A. thaliana*, and *O. sativa*. Homologous gene IDs were retrieved from NCBI (https://www.ncbi.nlm.nih.gov) or Phytozome v13 (https://phytozome-next.jgi.doe.gov). NLPs whose FRs were used in the LjNIN ($FR^{swap}$) are highlighted in bold. The tree was constructed by maximum likelihood (ML) method using IQ-TREE 2 (JTT+F+I+G4 substitution model). The percentages of replicate trees in which the associated taxa clustered together in the bootstrap test (1000 replicates) are shown next to the branches. Branch lengths are in the same units as those of the evolutionary distances. (**B**) Protein sequence alignment of FR of LjNIN and its closest homologous NLPs collected in (A). Highlighted letters on a gray background show the same amino acids as $FR^{NIN}$. Boldfaces on white background show amino acids with similar properties to those in $FR^{NIN}$. (**C**) Shoots (upper panels), ITs (middle panels) and nodules (lower panels) complemented by LjNIN proteins in which the FR was swapped with FRs shown in (B). LjNIN ($FR^{swap}$) proteins were expressed under *CE* (5 kb)-*LjNIN* promoter (3.3 kb) in hairy roots of *nin-9* mutants. Transgenic plants were inoculated with DsRED-labelled rhizobia for 28 d. Numbers indicate the frequency of plants shoot recovered (upper panels), forming ITs (middle panels) or pink nodules (lower panels) among all transgenic plants. Bars, upper panels = 1 cm, middle panels = 50 μm, lower panels = 2 mm.

**Table S1.**
**List of genes up-regulated by rhizobial inoculation in WT at 7 dai (compared to the expression level in WT at 0 dai ($log_2FC > 1$ and FDR < 0.05)) (4466 genes).**

**Table S2.**
**List of genes up-regulated by rhizobial inoculation in WT at 7 dai and decreased in ΔFR nin compared to in WT at 7 dai ($log_2FC < -1$ and FDR < 0.05)) (1165 genes).**

**Table S3.**
**List of genes up-regulated by rhizobial inoculation in WT at 7 dai and decreased in nin-9 compared to in WT at 7 dai ($log_2FC < -1$ and FDR < 0.05)) (1343 genes).**

**Table S4.**
**List of genes up-regulated by rhizobial inoculation in WT at 21 dai (compared to the expression level in WT at 0 dai ($log_2FC > 1$ and FDR < 0.05)) (6824 genes).**

**Table S5.**
**List of genes up-regulated by rhizobial inoculation in WT at 21 dai and decreased in ΔFR nin compared to in WT at 21 dai ($log_2FC < -1$ and FDR < 0.05)) (1979 genes).**

**Table S6.**
**List of genes up-regulated by rhizobial inoculation in WT at 21 dai and decreased in nin-9 compared to in WT at 21 dai ($log_2FC < -1$ and FDR < 0.05)) (2414 genes).**

**Table S7.**
**Primers used in this study**
